# Approximating carbon fixation - how important is the Calvin-Benson cycle steady-state assumption?

**DOI:** 10.1101/2022.11.18.517021

**Authors:** Marvin van Aalst, Oliver Ebenhöh, Berkley J. Walker

## Abstract

Plants use light energy to produce ATP and redox equivalents for metabolism. Since during the course of a day plants are exposed to constantly fluctuating light, the supply of ATP and redox equivalents is also fluctuating. Further, if the metabolism cannot use all of the supplied energy, the excess absorbed energy can damage the plant in the form of reactive oxygen species. It is thus reasonable to assume that the metabolism downstream of the energy supply is dynamic and as being capable of dampening sudden spikes in supply is advantageous, it is further reasonable to assume that the immediate downstream metabolism is flexible as well. A flexible metabolism exposed to a fluctuating input is unlikely to be in metabolic steady-state, yet a lot of mathematical models for carbon fixation assume one for the Calvin-Benson-Bassham (CBB) cycle. Here we present an analysis of the validity of this assumption by progressively simplifying an existing model of photosynthesis and carbon fixation.

## Introduction

The light reactions of photosynthesis must provide the chemical energy (ATP and NADPH) needed for CO_2_ fixation through the CBB cycle under rapidly fluctuating light availability. Light fluctuates across many time scales under natural conditions, greatly complicating the challenge plants have in harvesting sufficient light energy for optimal rates of carbon fixation [19, 3, 22, 16]. Plants possess a myriad of physiological responses to changing light intensity such as variable stomatal conductance, regulation of CBB enzymes and possibly even the activation of photorespiratory genes [4]. While these factors are critical for understanding the integrated response of net assimilation to fluctuating light, we have focused this investigation on dissecting out the important assumptions for modeling the CBB and photosynthetic electron transfer chain (pETC) using reaction kinetic models.

There have been many excellent metabolic models representing the CBB and associated pETC activity using various frameworks that can represent steadystate or dynamic behavior [2, 13, 18, 27, 12]. In this paper we use a metabolic definition of steady-state, specifically that under steady-state metabolite pool sizes are constant as well as all input and output fluxes. We use a corresponding metabolic definition for dynamic in that it is a condition where metabolite pool sizes and internal fluxes are changing. Note that this is a slightly different definition of a physiological dynamic model, which can represent components like the slow relaxation of non-photochemical quenching, changes in stomatal conductance and rubisco activation state [23]. In this work we ask how important it is to consider the dynamic response of the CBB during light fluctuations and how well the commonly applied steady-state assumption can represent metabolism.

The fundamental aim of mathematical modeling is to enhance understanding of the system studied, as B.D. Hahn already noted in 1993 [8]. Thirty years later much effort is still being put into building ever more complicated models with high fidelity, which are increasingly difficult to understand. Notably, Hahn considered models with 17-31 non-linear differential equations to be large while nowadays genome-scale modeling techniques are frequently employed, which can contain thousands of reactions [15, 6, 7, 11]. To revisit the spirit of trying to understand the key features of a system we employ model reduction of a previously published model of the pETC and the CBB cycle to elucidate the main processes controlling carbon fixation in C3 plants under dynamic conditions [21]. The resulting simplified models can serve as robust alternatives in situations where few parameters are known and the research question is only concerned with carbon fixation rate, as their predictions are in very good agreement with the predictions of the original complex model. The three consecutive reductions we employ are first replacing the dynamic system behavior with a steady-state approximation, then replacing the steady-state system with a polynomial fitted to the predicted carbon dioxide fixation rate and lastly reducing the amount of data that is fed into the model. We find that a good prediction of carbon fixation rate requires a remarkably low amount of model fidelity as well as data, while prediction of other fluxes requires a much better description of the underlying biochemistry.

## Results

To determine the required fidelity to represent the CBB cycle, pETC and photo-protective mechanisms, we first compared simulations of a detailed ordinary differential equation (ODE) model with and without a simplifying metabolic steady-state assumption [21, 13]. For this first simplification we removed the dynamic change of metabolite concentrations by pre-calculating steady-state fluxes of an ODE model in response to realistic field conditions using incoming photosynthetically active photon flux density (PPFD) measured in one minute intervals at a National Ecological Observatory Network site in Washington state, USA [17]. We then compared the simulated rate of carbon fixation of the dynamic and the steadystate model for a 6 hour window from a typical summer day (shown in Figure 1). There is generally good agreement between the simulation of carbon fixation (ribulose-1,5-bisphosphate carboxylase-oxygenase (rubisco) flux) between the two approaches. While the relative error in short periods can exceed 50 %, the total error of predicting the rubisco flux is 0.49 %.

**Figure 1.**
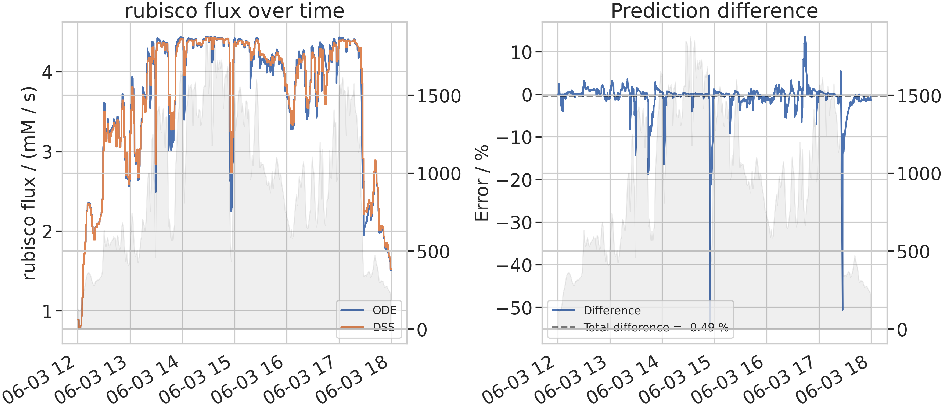
Comparison of ODE and steady-state approximated model predictions over a dynamic light signal (grey area). The left subplot shows the rubisco flux predicted by the ODE and the steady-state approximated model respectively over a time course of 6 hours. The right subplot shows the error of the steady-state approximated model rubisco flux predictions relative to ODE model predictions.

To understand why the steady-state model behaves similarly to the dynamic model with regard to total carbon fixation despite the lack of dynamic interaction we investigated how well other fluxes are predicted and how much the concentrations change over the course of the experiment. Shown in Figure 2 is the absolute total difference between the simulations of key CBB, pETC and acclimation fluxes and the relative standard deviation (coefficient of variation) of representative concentrations in the dynamic model. The remaining fluxes and concentrations are displayed in supplementary figures S1 and S2 respectively and the relative standard deviation for the fluxes in supplementary Figure S3. In contrast to the the small difference between the rubisco flux in the simulations, the difference between rapid acclimation response mechanisms such as the xanthophyll cycle (zeaxanthin epoxidase) or reactions of the water-water cycle (glutathione reductase) is between 20 % and 50 %. Similarly, the relative standard deviation of the rubisco substrate ribulose-1,5-bisphosphate (RuBP) is comparatively small, varying 15 % of its mean value compared to the large relative standard deviation of some metabolites that take part in the rapid acclimation response, e.g. glutathione-disulfide (GSSG) and dehydroascorbate (DHA) which vary nearly up to 350 % of their mean value. Notably, other CBB cycle intermediates like sedoheptulose-1-7-bisphosphate (SBP) and sedoheptulose-7-phosphate (S7P) vary up to 50 % while the difference between the flux predictions of sedoheptulose-bisphosphatase (SBPase) is comparable to the one of rubisco.

**Figure 2.**
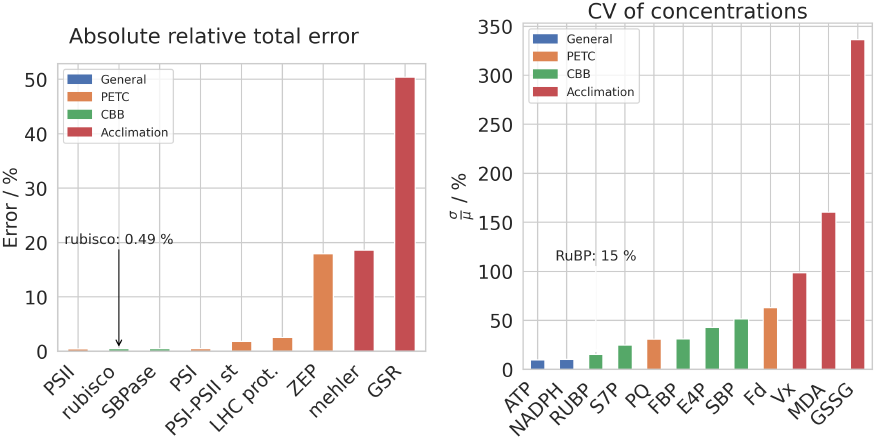
Left: absolute total error of steady-state approximated model predictions compared to ODE model predictions. Right: relative standard deviation (coefficient of variation) of ODE model predicted metabolite concentrations over a course of a 6-hour experiment.

To understand how alternative parameterizations of the model that lead to similar carbon fixation rates would effect internal fluxes of the CBB cycle we generated 100,000 sets of randomly perturbed kinetic parameters. For these parameter-sets we calculated the steady state flux and then selected the ones for which the carbon fixation flux was within 1 % of the original model, which was the case for 321 of them. Figure 3 shows a box-plot of simulated CBB cycle fluxes of the selected parameter sets relative to the dynamic model, the remaining fluxes are shown in supplementary Figure S7. Some reactions of the CBB cycle must vary less than 10 % to ensure a similar flux, while e.g. triose-phosphate isomerase (TPI), fructose-1,6-bisphophatase (FBPase) and fructose-bisphosphate aldolase (ALDOA) can vary more than 30 %. Reactions related to triose phosphate export and storage can vary upwards of 100 %.

**Figure 3.**
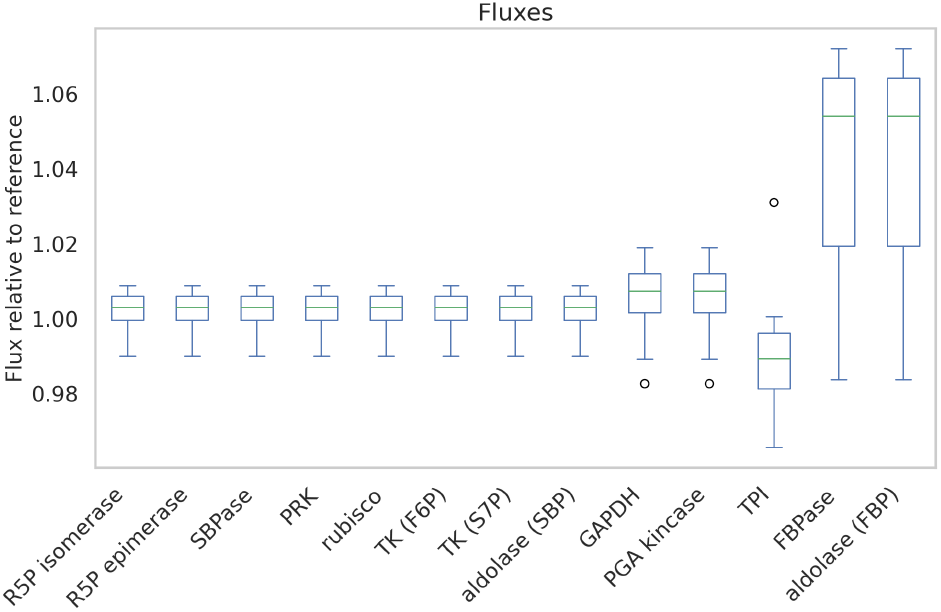
Box-plot of CBB cycle fluxes in model collection with randomly perturbed kinetic parameters relative to the fluxes in model with reference parameters.

Given the stability of the RuBP concentration during our simulations across irradiances and the insensitivity of rubisco flux to parameterization of many CBB model reactions, we hypothesized that it is possible to vastly simplify the CBB cycle and still get a reasonable prediction. To describe the carbon fixation rate as being only dependent on irradiance, we replaced the explicit CBB model with a 4th degree polynomial function which we fitted over simulated carbon fixation fluxes. The agreement between these two models was very good, with a very small root-mean-square error (RMSE) of 0.02 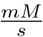 (shown in supplementary Figure S4). When simulated over the same data shown in Figure 1, the difference between the total rubisco flux of the polynomial model and the dynamic model ODE fluxes is 0.64 % and thus comparable to the difference of 0.49 % between the steady-state model and the dynamic model (shown in supplementary Figure S5).

So far we have used high temporal resolution data in our simulations, but data availability, lack of computational power or technical difficulties due to multiscale modeling can require the use of low temporal resolution data. A common practice to reduce the temporal resolution is to simply average the PPFD values over longer time periods. In our experiment, an increase from one minute to 60 minute steps increased the difference of carbon fixation from 0.49 % to roughly 7 % (see supplementary Figure S10).

While an increase in error with fewer data was to be expected, we hypothesized that the simulation can be improved substantially by using an alternative averaging approach. If carbon assimilation responded linearly to irradiance, carbon assimilation over the average irradiance of a time period would equal the average of carbon assimilation over that time period. In other words, irradiance could be averaged over a time period with fluctuating light and simulate an equal total amount of carbon fixation as the fluctuating light period. However, as carbon fixation responds non-linearly to irradiance, the simulated carbon fixation of the average irradiance does not lead to the average of the simulated carbon fixation over fluctuating light. In the case of the saturating response of carbon fixation to irradiance, simple averaging would lead to over-estimates over periods of fluctuating irradiance.

To produce representative PPFD values that avoid the bias of simple averaging over each time period, we clipped the input data by capping saturating PPFD values to reduce their effect before calculating the mean. As this approach led to promising results with our single-day data we expanded the analysis to a representative day for each month of the entire year, and then simulated the carbon fixation rate of this day for each month with one minute steps for the ODE model and 60 minute steps for the polynomial model.

For the polynomial model we included both the prediction with the mean of the raw data and the mean of the clipped data. Figure 4 shows the total error of the rubisco flux prediction of the polynomial model relative to the ODE model per month. As indicated in the figure legend, the total error per year of the polynomial model relatives to the ODE model is 4.0 % for raw data and 0.1 % for data clipped at a PPFD of 1000. The absolute of the error of the polynomial model for October to February is between 5 and 15 %, while it is lower for the summer months (except June), however if the data is clipped at a PPFD of 1000 there is an improved fit relative to the polynomial model for the winter months.

**Figure 4.**
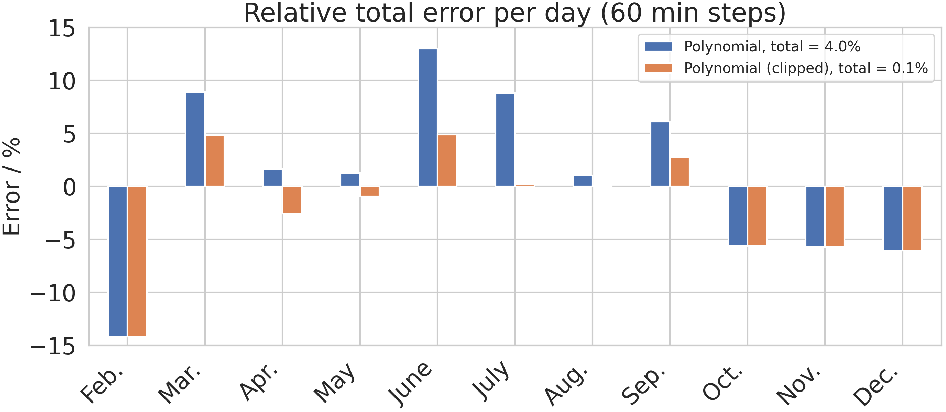
Total prediction error of carbon fixation of the steady-state approximated model (60 minute resolution) and the polynomial model (60 minute resolution) relative to the ODE model (1 minute resolution). Results are shown for both the mean PPFD value of the raw data and the mean value of the data clipped at either PPFD 900 (steady-state approximated model) or 1000 (polynomial model).

## Discussion & Conclusions

Our simulations reveal that accounting for the dynamic change of metabolite concentrations under fluctuating light results in only small differences in the predicted carbon fixation rate relative to assuming a metabolic steady state in a combined model including the CBB and light reactions. Specifically, in Figure 1 we compared predictions of an ODE model with a simplified version in which we removed the dynamic change of metabolite concentrations and instead precalculated the steady-state fluxes for a range of PPFD inputs. Despite the rapid light fluctuations in the input data, our results show that the predicted total carbon fixation essentially stays the same, even though the difference of the prediction of the dynamic state can exceed 50 % for some time steps. This means that for the purpose of predicting total carbon fixation, our ODE model of carbon fixation can be greatly simplified using steady-state assumptions with little difference - saving time and energy by avoiding computationally costly numerical integration. We expect this to hold for other ODE models of biological systems that are structured similarly. Next we investigated how such a simplified model, which does a mechanistically poor job at representing the underlying biochemistry, does a good job at representing carbon fixation. For this we turned to other biochemical predictions of the model.

While total carbon fixation was predicted well in the steady-state model, the fluxes of rapid acclimation mechanisms (e.g. violaxanthin deepoxidase) were predicted poorly, with relative errors exceeding 50 % (see Figure 2). These observations can be explained by the amount of dynamic change in concentration over time of specific metabolite pools that directly influence carbon fixation. For example, the RuBP pool is comparatively stable (varying 15 % of its mean concentration), while the GSSG pool can vary over 350 %of its mean concentration (see Figure 2). Since the RuBP pool comprises the only dynamic substrate for carbon fixation in the original model it has the largest effect on rates of rubisco carboxylation, reflected by the small relative error in this reaction of 0.49 %. The higher errors of acclimation and pETC intermediates in contrast contribute very little to the error of rubisco carboxylation, as they mostly depend on prior perturbations of the system but always tend towards stabilizing the system towards the respective steady state. Notably, other CBB cycle intermediates like SBP can also show a variation of more than 50 % of their mean concentration, but these errors are not reflected in the downstream concentrations of RuBP, resulting in less differences in rates of rubisco carboxylation.

One explanation for the different variation over time in metabolite pool sizes is that multiple flux distributions can lead to the same carbon fixation rate, thus allowing a wider range of concentrations for certain CBB cycle intermediates. This buffering effect can be seen in Figure 3, where in models with perturbed kinetic parameters but similar rates of carbon fixation some reactions like FBPase can vary more than a third of their reference flux. This suggests that the CBB cycle is structured such that temporary changes in illumination are buffered by the cycle for RuBP concentration to remain relatively constant, resulting in similar rates of carbon fixation, effectively working as a low-pass filter (see supplementary figures S8 and S13 and supplementary Figure S16 to Figure S45). Due to this stability, carbon fixation can be accurately simulated, even if the underlying metabolism is not. While the stability of RuBP is advantageous for downstream metabolism this also implies that alternative energy sinks like quenching mechanisms and the water-water cycle (WWC) are required to dissipate further excess energy. The reactions with the most variation relate to sucrose and starch partitioning (e.g. FBPase, FBP aldolase and TPI), suggesting that while carbon fixation is simulated accurately, downstream carbon partitioning may not be. These findings suggest that carbon fixation can be simulated accurately in an even more simplified CBB cycle, providing down-stream carbon partitioning is not of interest. Our findings that an explicit model of the CBB cycle can be replaced with a simple polynomial model with little sacrifice, see Figure 4, are in line with many crop-systems models that represent net carbon assimilation using a simple radiation use efficiency (e.g.

The discussion above highlights that metabolic models of the CBB cycle can produce similar rates of carbon fixation despite mechanistically curedely simplified assumptions of the underlying biochemistry. This finding indicates that care is needed when validating a model using only carbon fixation rates, since any model that keeps RuBP concentration stable can lead to realistic carbon fixation rates. Possible improvements include using predictions of other metabolite pools ([2]) or fluorescence data if the model also contains the pETC ([21]). Further work can test the ability to make steady-state assumptions with more complex models of carbon assimilation that include for example photorespiration, CO2 as a dynamic variable or dynamic temperature. Photorespiration may present an interesting case since large pools of glycine accumulate during photorespiratory induction, effectively decreasing relative rates of glycine decarboxylation (and increasing net carbon exchange) during this transient by up to 40 % [5].

We show that temporally high-resolution PPFD data is not required to give a good prediction of total carbon fixation as reducing the amount of data by 60-fold still led to an prediction error of less than 1 % if over modified averaging approach is used, see Figure 4. The ability to properly aggregate data is important, since often all data needed to produce and validate a model are not available on the same time resolutions. For example, in canopy scale predictions, irradiance values are available at time scales of seconds, while eddy covariance data is usually presented over 30 minute time steps. Our modeling indicates that clipping saturating values before averaging improved prediction error by ≈14-fold, see Figure 4. While these findings demonstrate the value of clipping saturating values before averaging, considering the saturating kinetics of carbon fixation to PPFD, this method could be valuable for any high-resolution data which needs to be averaged over a time-step in a process with saturating kinetics.

## Methods

We performed our analyses in Python 3.10, utilising the common packages NumPy, pandas, SciPy and Matplotlib for general data analysis as well as modelbase and Assimulo for building and integrating the ODE model [20, 9, 14, 25, 10, 24, 1]. All code used to generate the publication results and figures is publicly available on our GitLab repository https://gitlab.com/qtb-hhu/photosynthesis-task-force/2022-how-realistic-is-the-cbb-ssa. We obtained field observation PPFD data measured in one minute intervals in Washington state (latitude 45.790835, longitude −121.933788) by the National Ecological Observatory Network (NEON) for which we identified which day had the highest data coverage for each month, which was the 25th, and the ODE model from Saadat 2021 [17, 21]. Due to model instabilities for PPFD values below 30, we clamped the minimum of the data to 30.

### Steady-state model

We calculated the steady state fluxes of the ODE model for PPFD values between the minimal and maximal values found in the dataset with step size 1. Then we simulated the model by looking up the steady state flux of the PPFD value rounded to the nearest integer.

### Approximations

We fitted the *V*_max_ and *K_m_* parameters of the Michaelis-Menten function using the SciPy `minimize` function and the L-BFGS-B algorithm [25, 26]. The polynomial fit was performed using NumPy’s `polyfit` function [9].

## Funding

This work was funded by the Deutsche Forschungsgemeinschaft (DFG) under Germany’s Excellence Strategy EXC 2048/1, Project ID: 390686111 (O.E.), EU’s Horizon 2020 research and innovation programme under the Grant Agreement 862087 (M.v.A.) and also supported by the U.S. Department of Energy Office of Science, Basic Energy Sciences under Award DE-FG02-91ER20021 (B.J.W.).

## Notes

### Competing Interest Statement

The authors have declared no competing interest.

### Summary of Updates

Funding information updated

