## supplementaries for "Approximating carbon fixation - how important is the Calvin-Benson cycle steady-state assumption?"

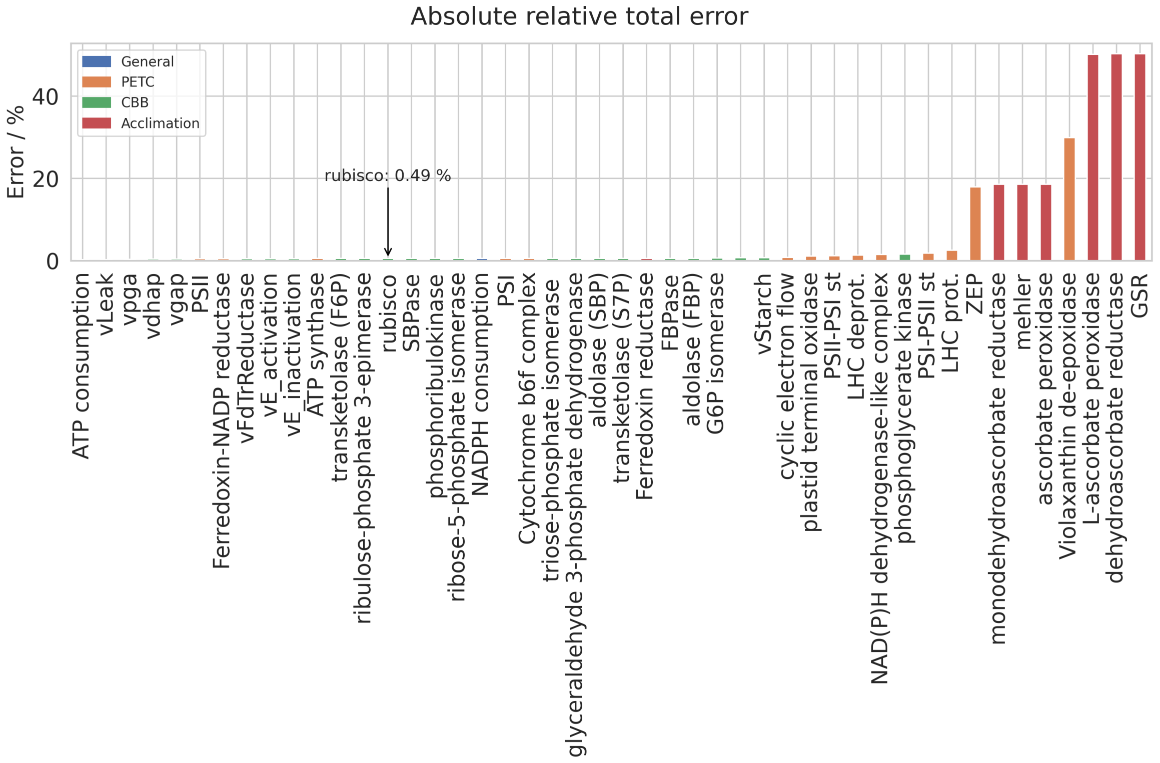


Figure S1: Absolute total error of flux predictions of the simplified model relative to the ordinary differential equation (ODE) model.

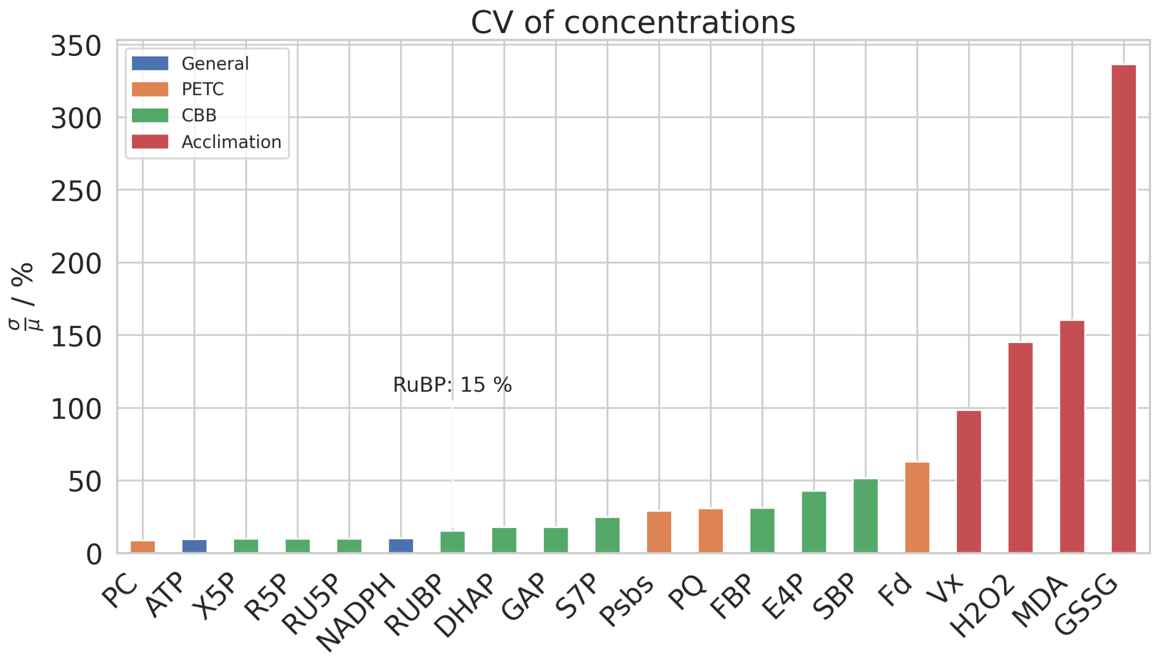


Figure S2: Relative standard deviation (coefficient of variation) of concentrations predicted by the ODE model over a course of a 6-hour experiment.

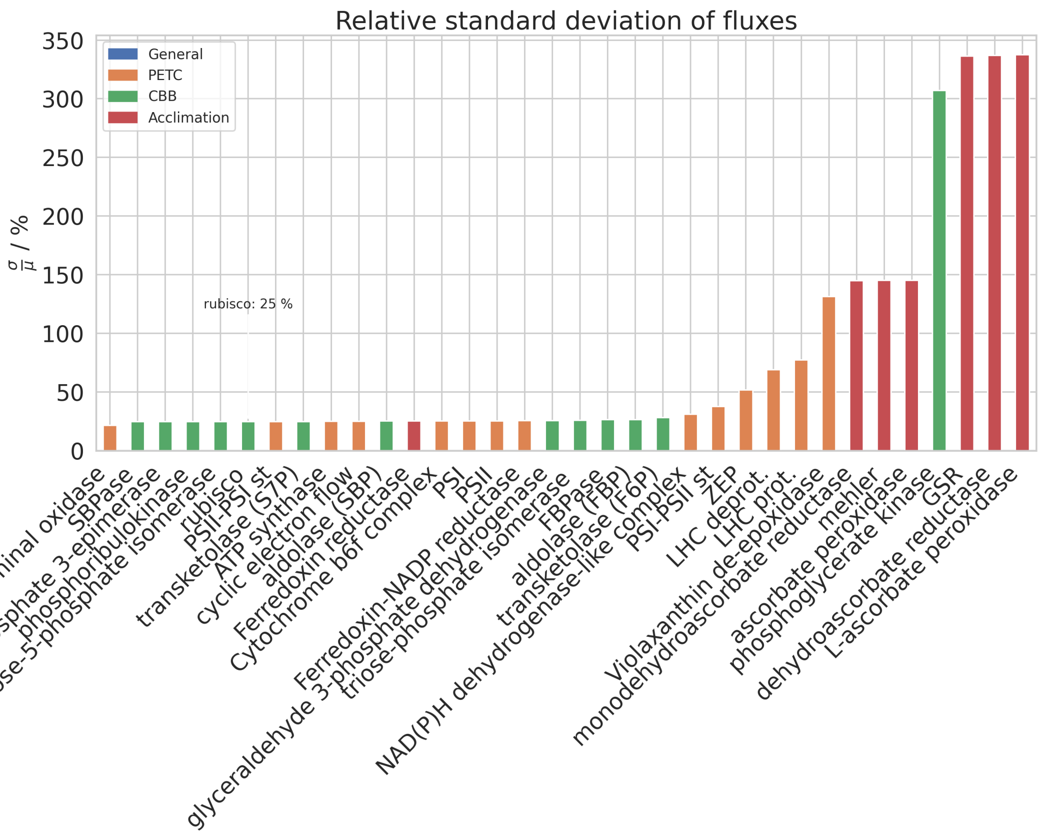


Figure S3: Relative standard deviation (coefficient of variation) of fluxes predicted by the ODE model over a course of a 6-hour experiment.

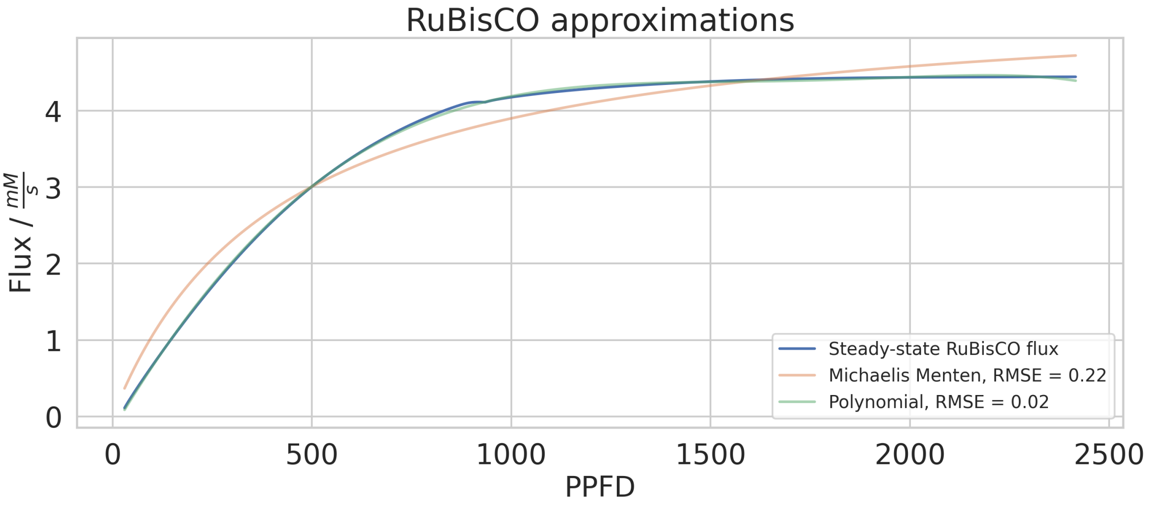


Figure S4: Steady-state scan of ribulose-1,5-bisphosphate carboxylase-oxygenase (rubisco) flux in the ODE model over a range of photosynthetically active photon flux density (PPFD) values with both a Michaelis-Menten and polynomial function fitted to the obtained data. The Michaelis-Menten function has a root-mean-square error (RMSE) of 0.22 $\frac{mM}{s}$, while the polynomial function has a RMSE of 0.02 $\frac{mM}{s}$.

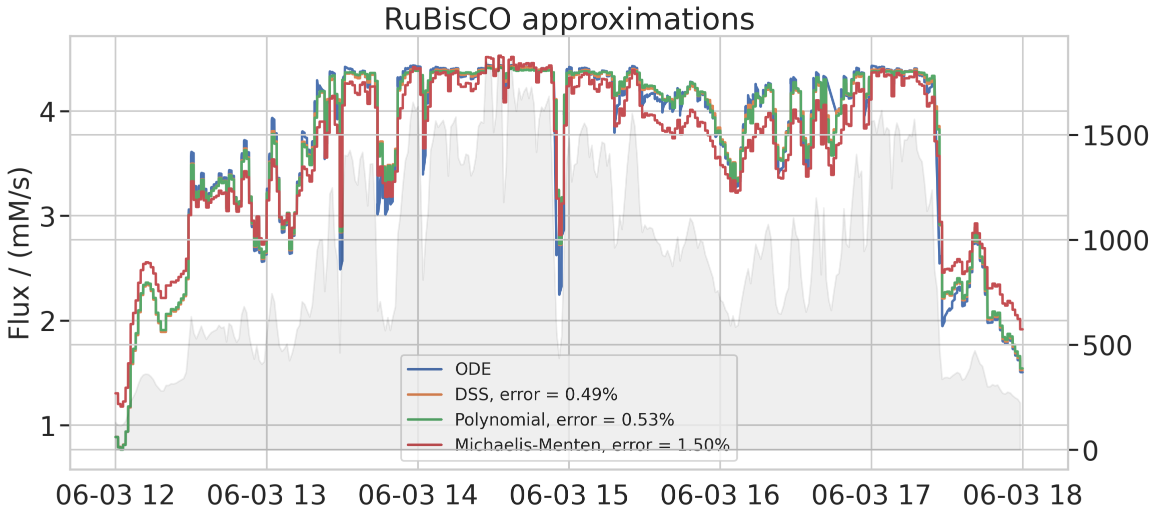


Figure S5: Prediction of rubisco flux by the ODE model, steady-state simplified model and polynomial model over a six hour time window. The legend shows the absolute total error of the flux relative to the ODE model.

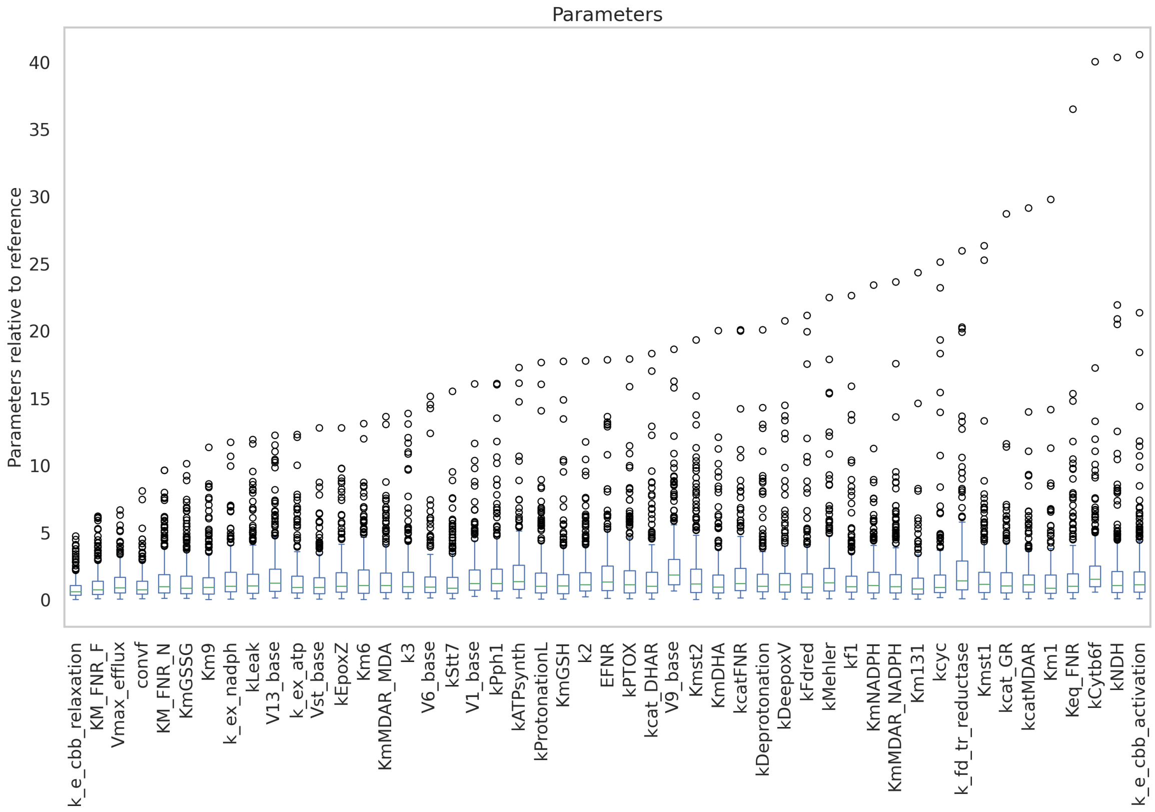


Figure S6: Distribution of parameter values relative to the parameter values of the original model that in respective combinations lead to a carbon fixation rate within 1 % of the original model.

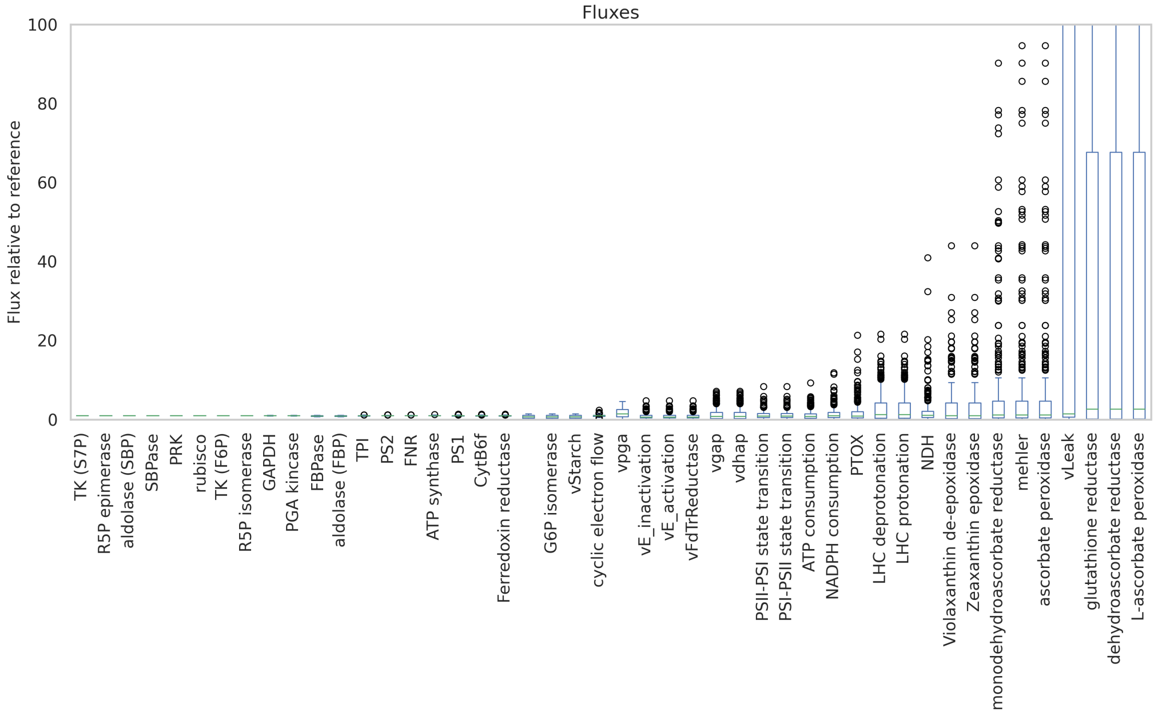


Figure S7: Distribution of fluxes relative to the fluxes of the original model that in respective combinations lead to a carbon fixation rate within 1 % of the original model.

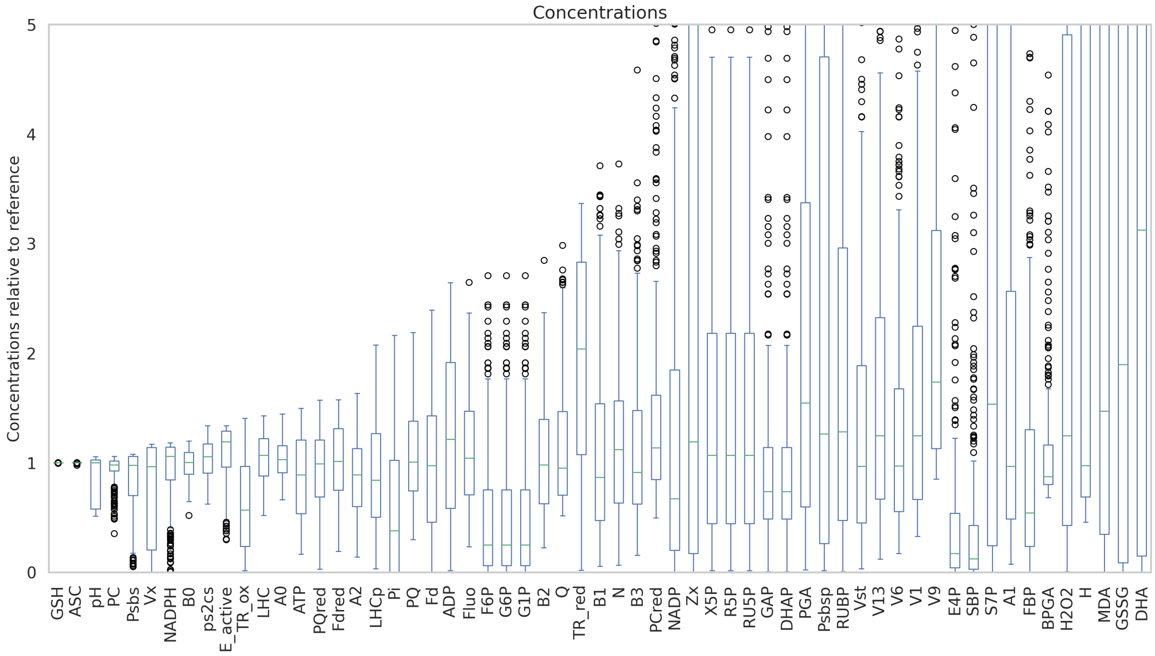


Figure S8: Distribution of concentrations relative to the concentrations of the original model that were produced by fluxes that lead to a carbon fixation rate within 1 % of the original model.

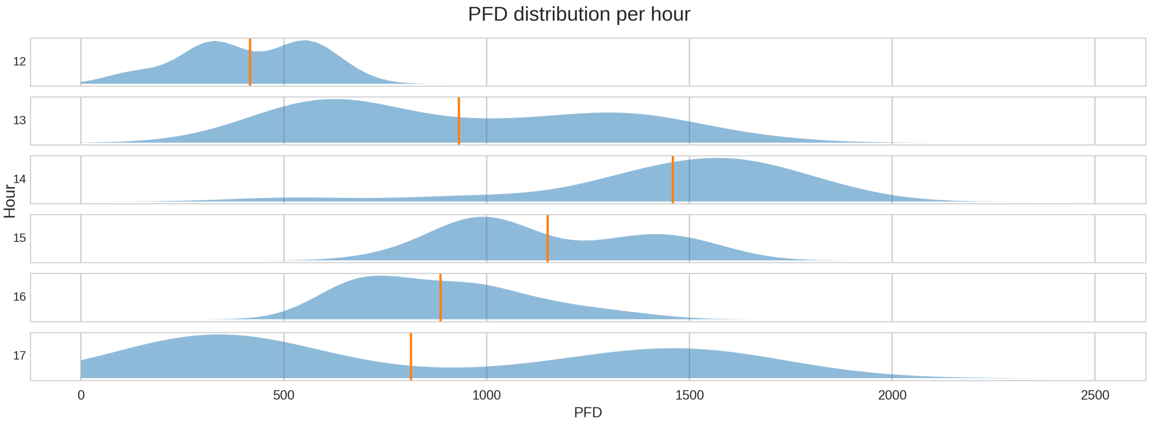


Figure S9: PPFD distribution per hour of day for a typical summer day.

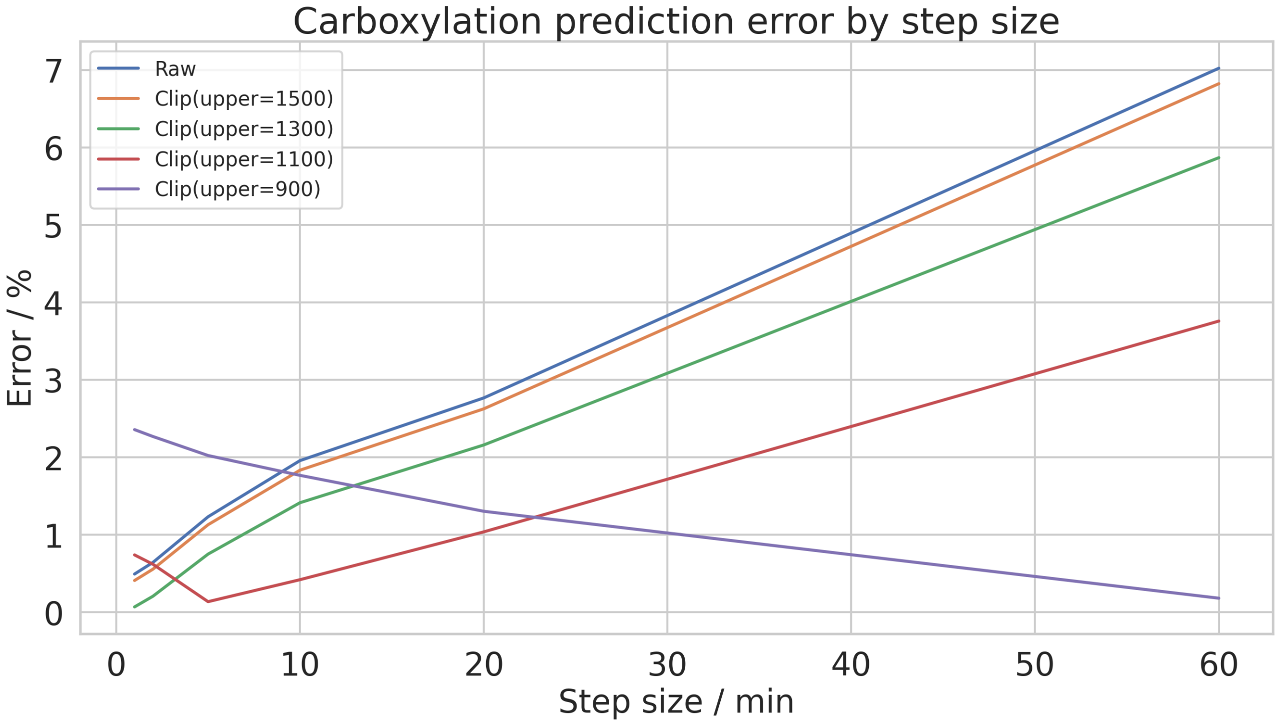


Figure S10: Error of predicted carboxylation flux of the M2 relative to the ODE model depending on step size and clipping point for PPFD values.

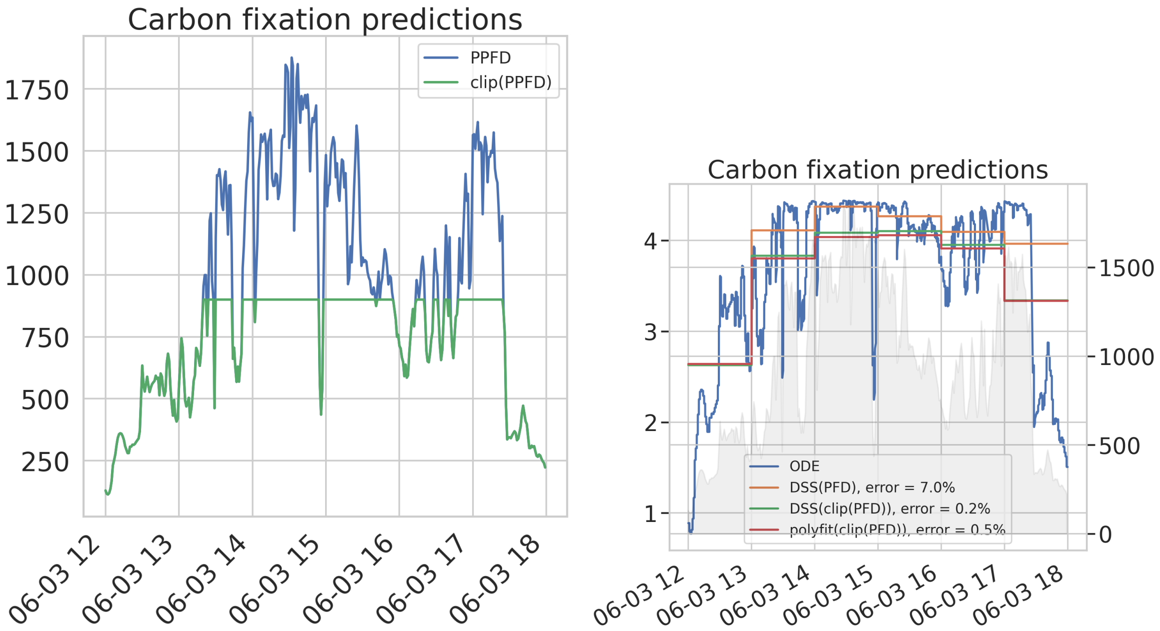


Figure S11: Clipping of the PPFD input signal on left-hand side graph and prediction of rubisco flux for either un-clipped data (M1) or clipped data (M2 and M3) on right-hand side graph.

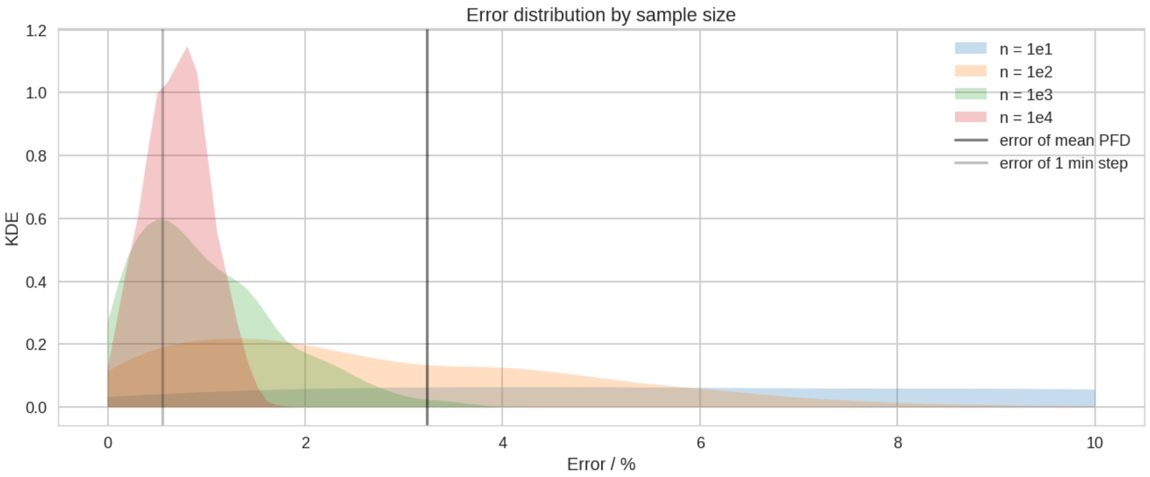


Figure S12: Error of M2 carbon fixation flux prediction relative to the ODE model by utilizing either the mean PPFD value of a 60 minute sample with one minute time steps, the mean flux of 60 PPFD values or the mean flux of random samples with different sample sizes.

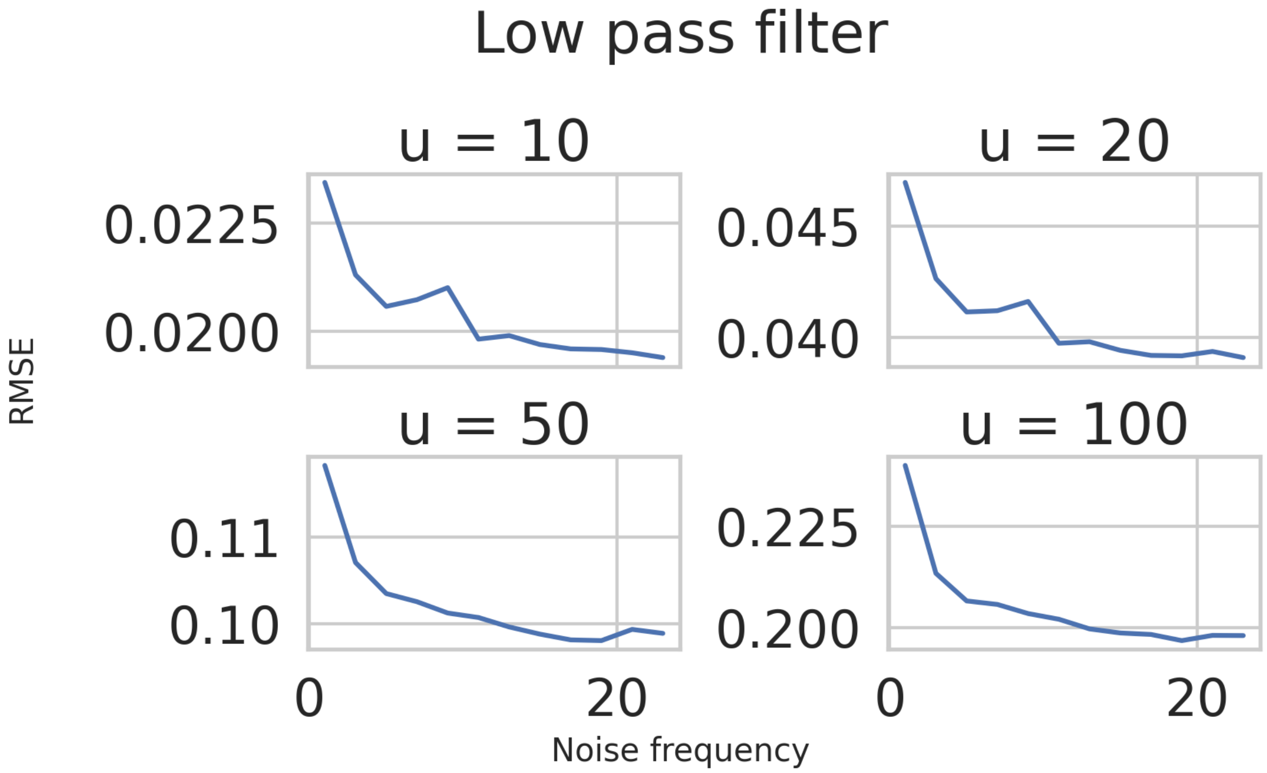


Figure S13: Difference between the carbon fixation prediction of the ODE for different PPFD inputs. The base input is a sine wave centered around PPFD 700 $\frac{\mu mol}{s \cdot m^2}$ with an amplitude of 300 $\frac{\mu mol}{s \cdot m^2}$ and frequency of 1 / hour, onto which a noise signal with an amplitude between 10 and 100 $\frac{\mu mol}{s \cdot m^2}$ and a frequency between 0 and 23 / hour is added.

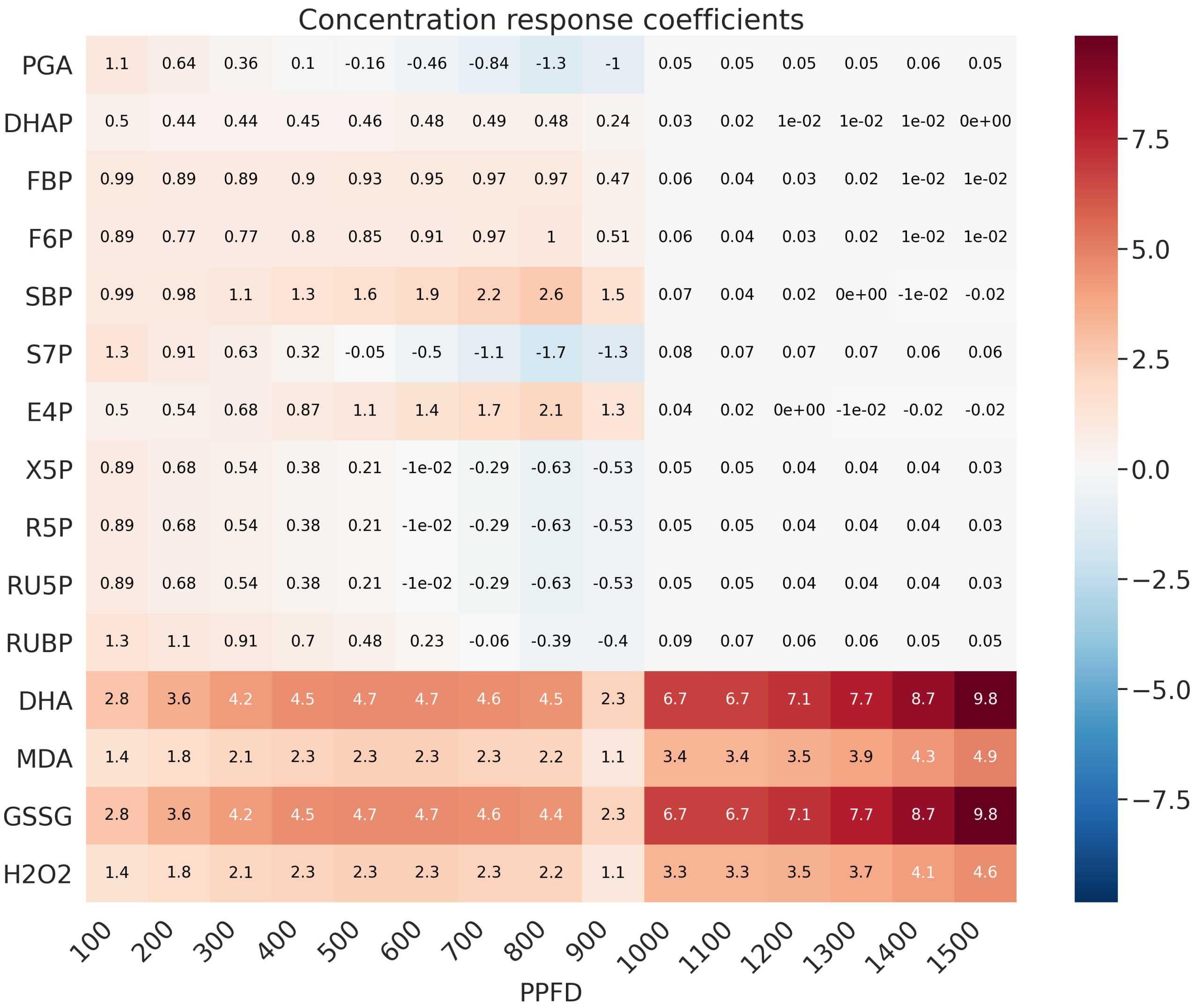


Figure S14: Concentration response coefficients to small changes in PPFD at different illumination levels.

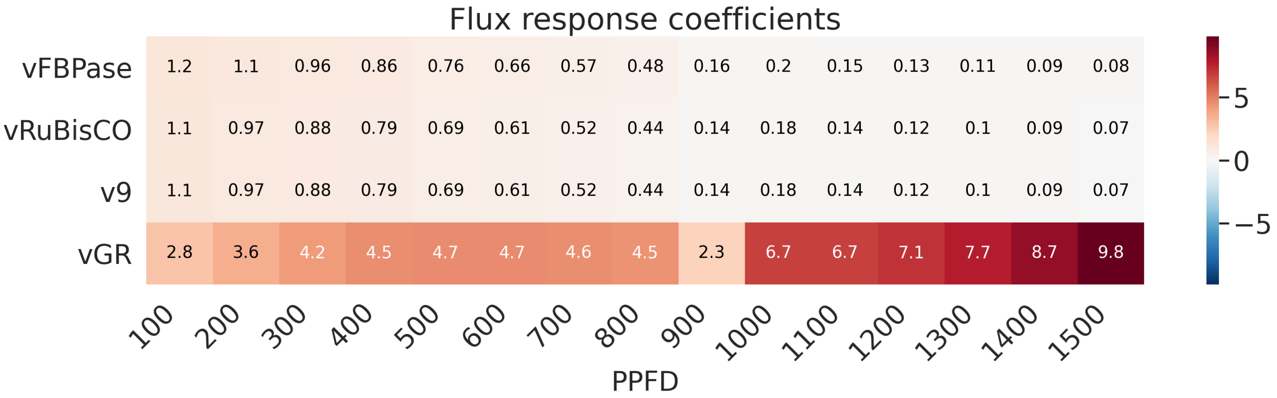


Figure S15: Flux response coefficients to small changes in PPFD at different illumination levels.

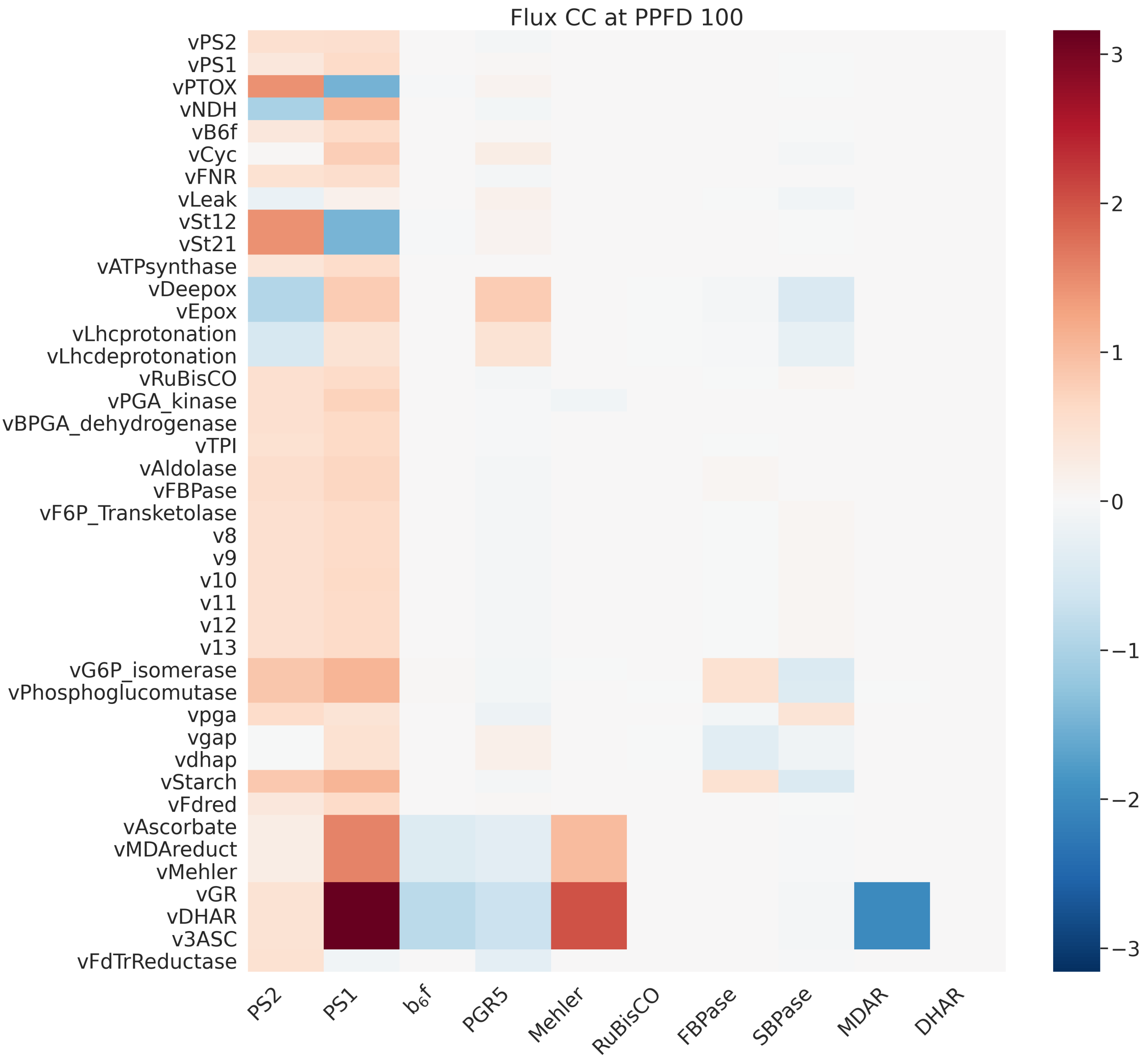


Figure S16: Flux control coefficients of representative reactions of photosynthesis at specified PPFD.

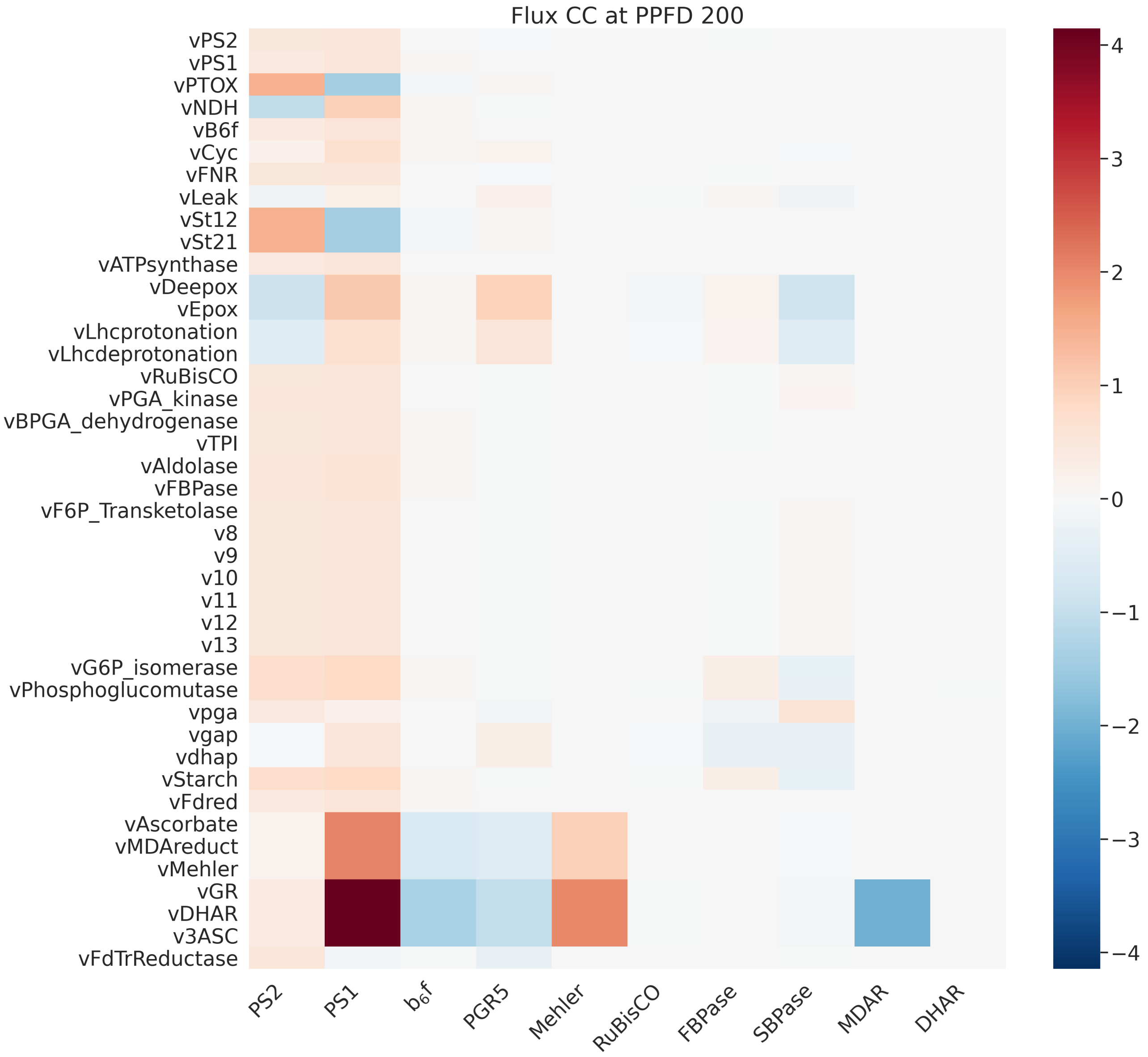


Figure S17: Flux control coefficients of representative reactions of photosynthesis at specified PPFD.

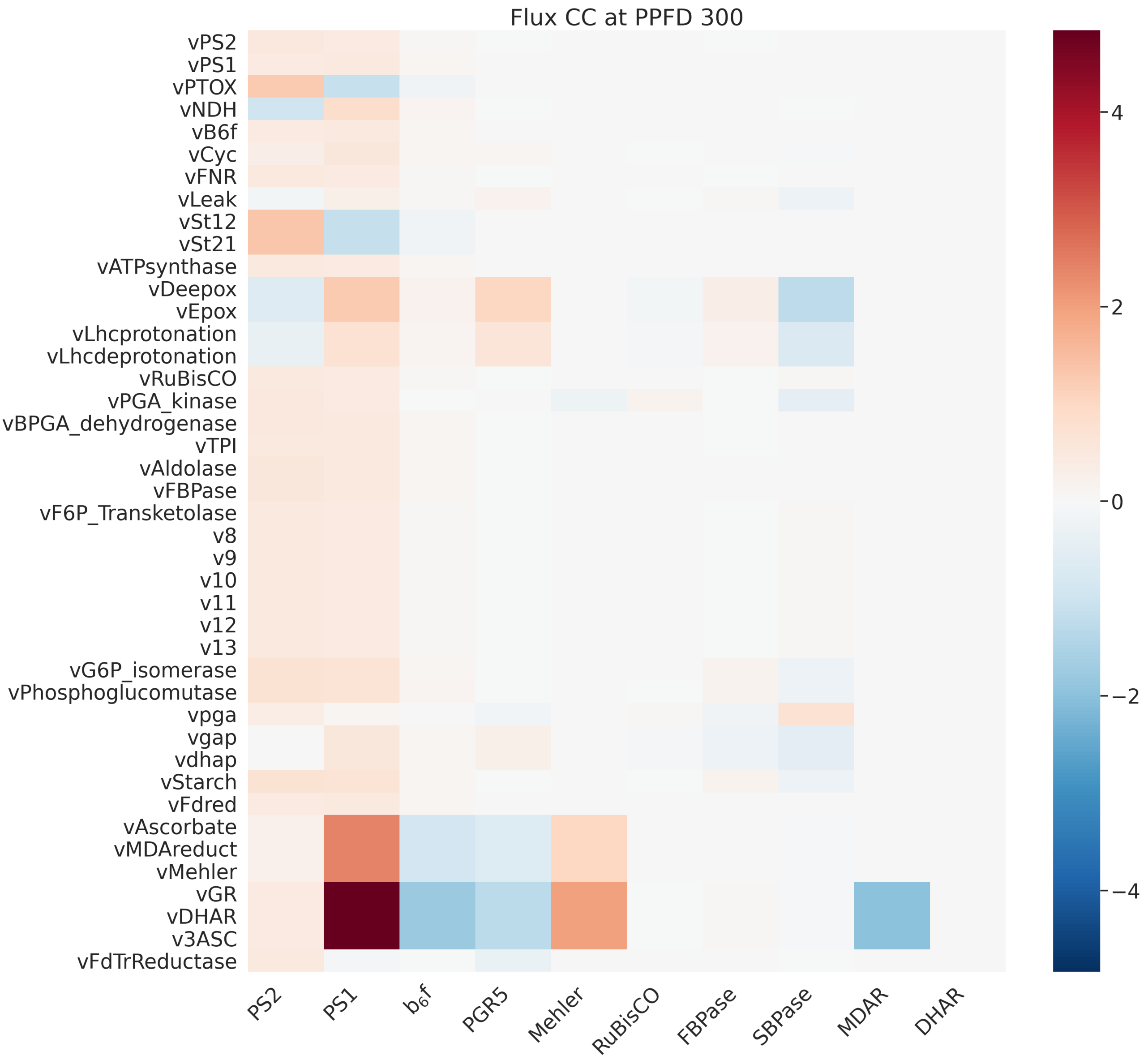


Figure S18: Flux control coefficients of representative reactions of photosynthesis at specified PPFD.

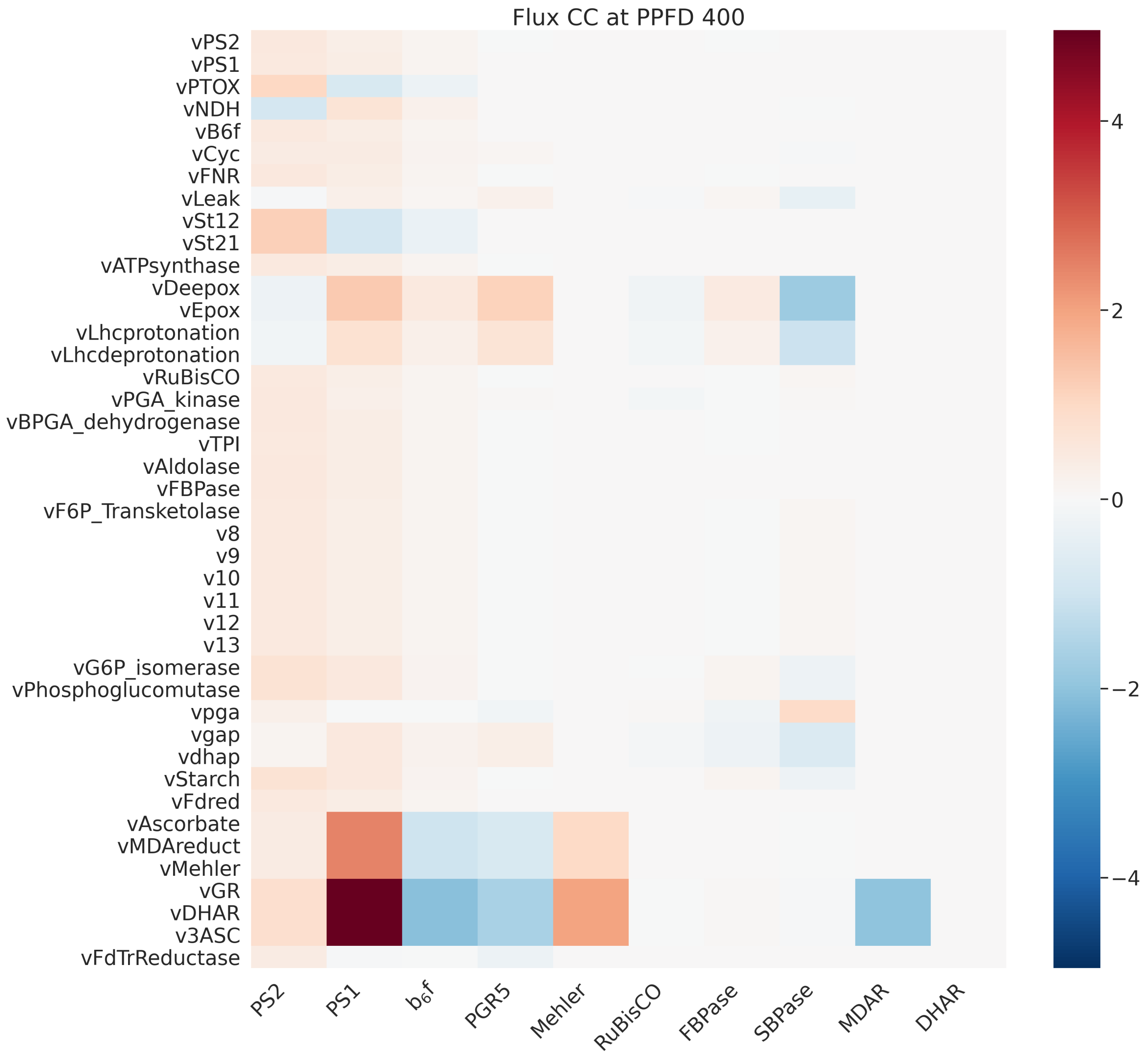


Figure S19: Flux control coefficients of representative reactions of photosynthesis at specified PPFD.

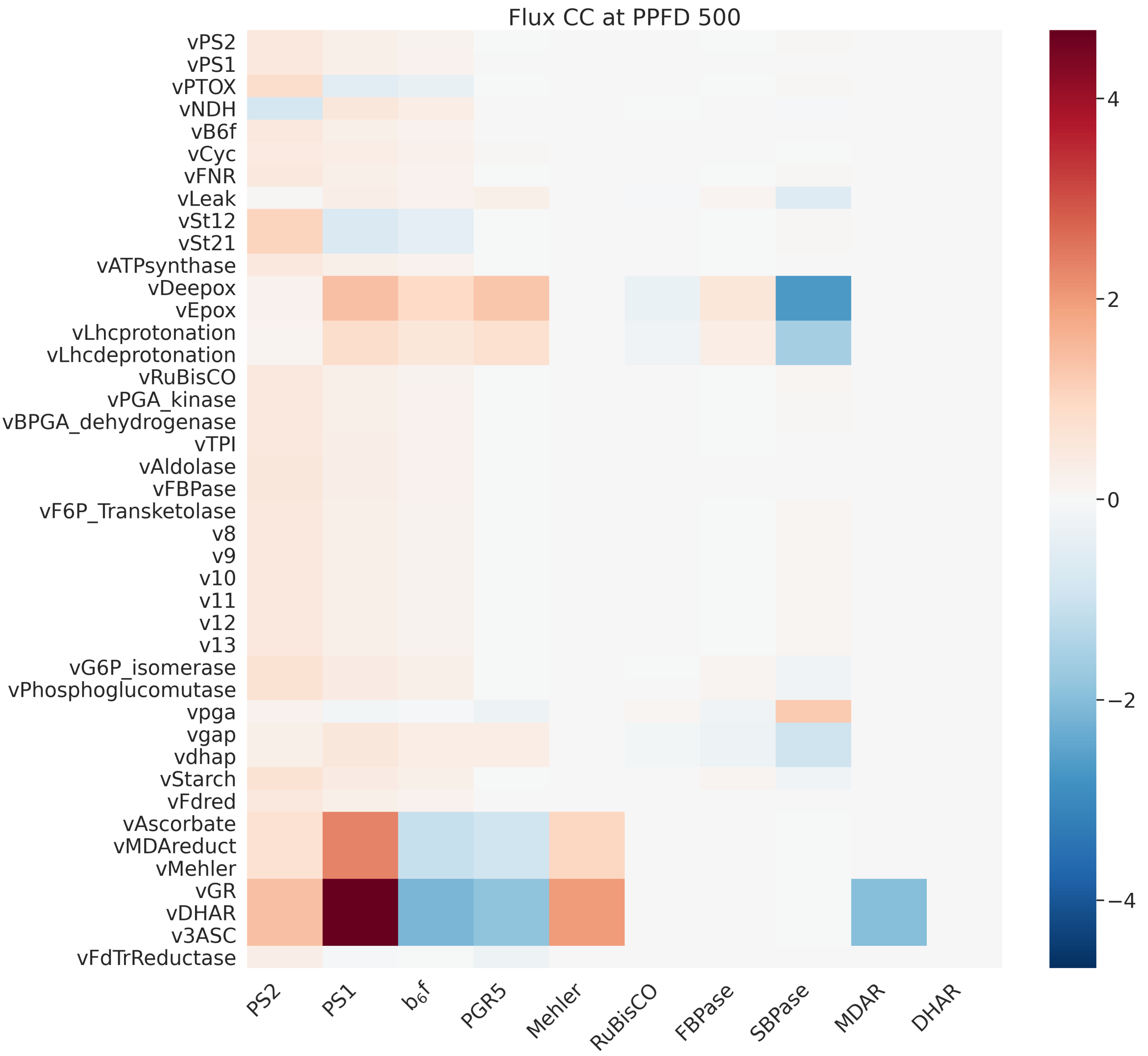


Figure S20: Flux control coefficients of representative reactions of photosynthesis at specified PPFD.

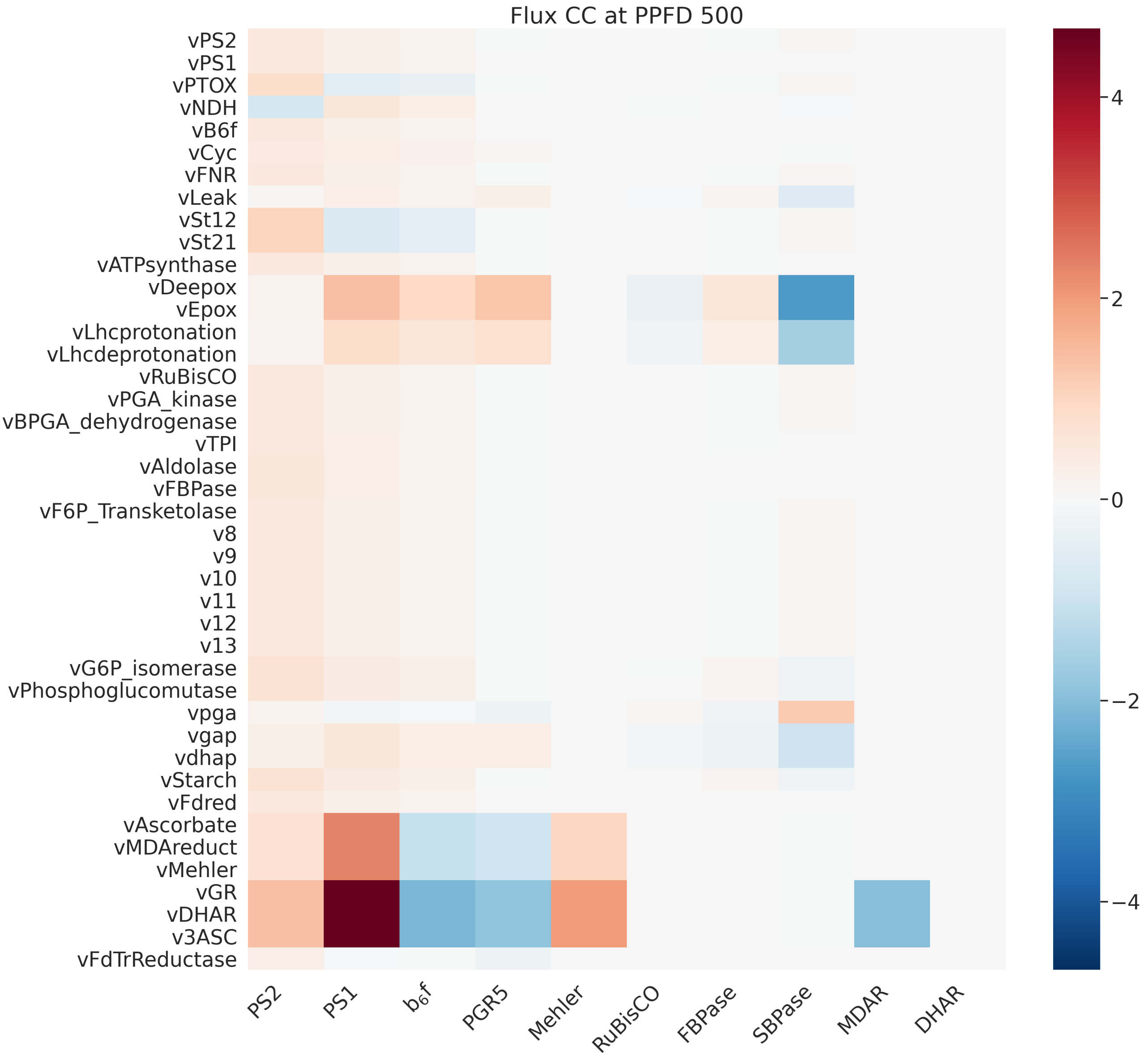


Figure S21: Flux control coefficients of representative reactions of photosynthesis at specified PPFD.

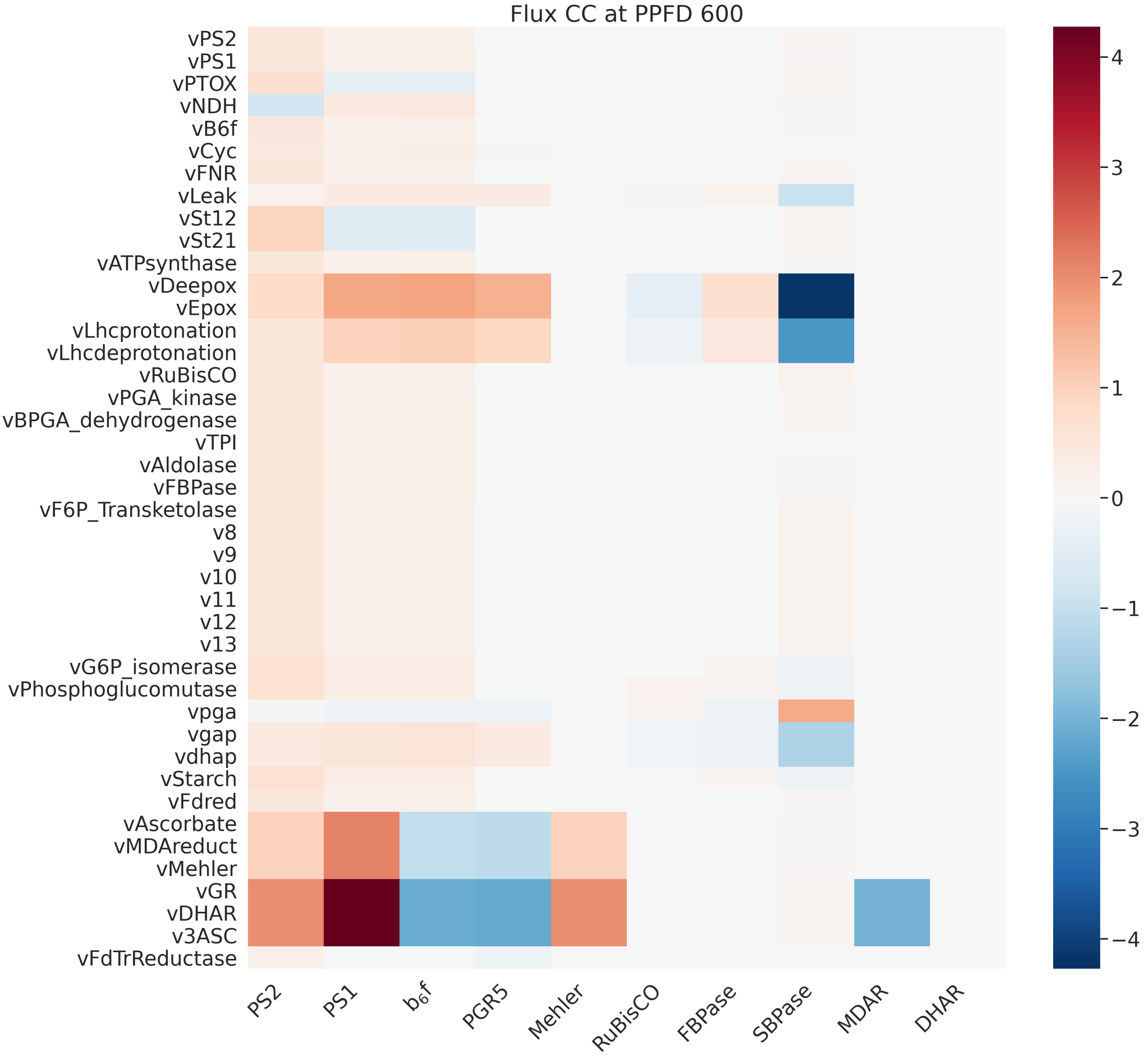


Figure S22: Flux control coefficients of representative reactions of photosynthesis at specified PPFD.

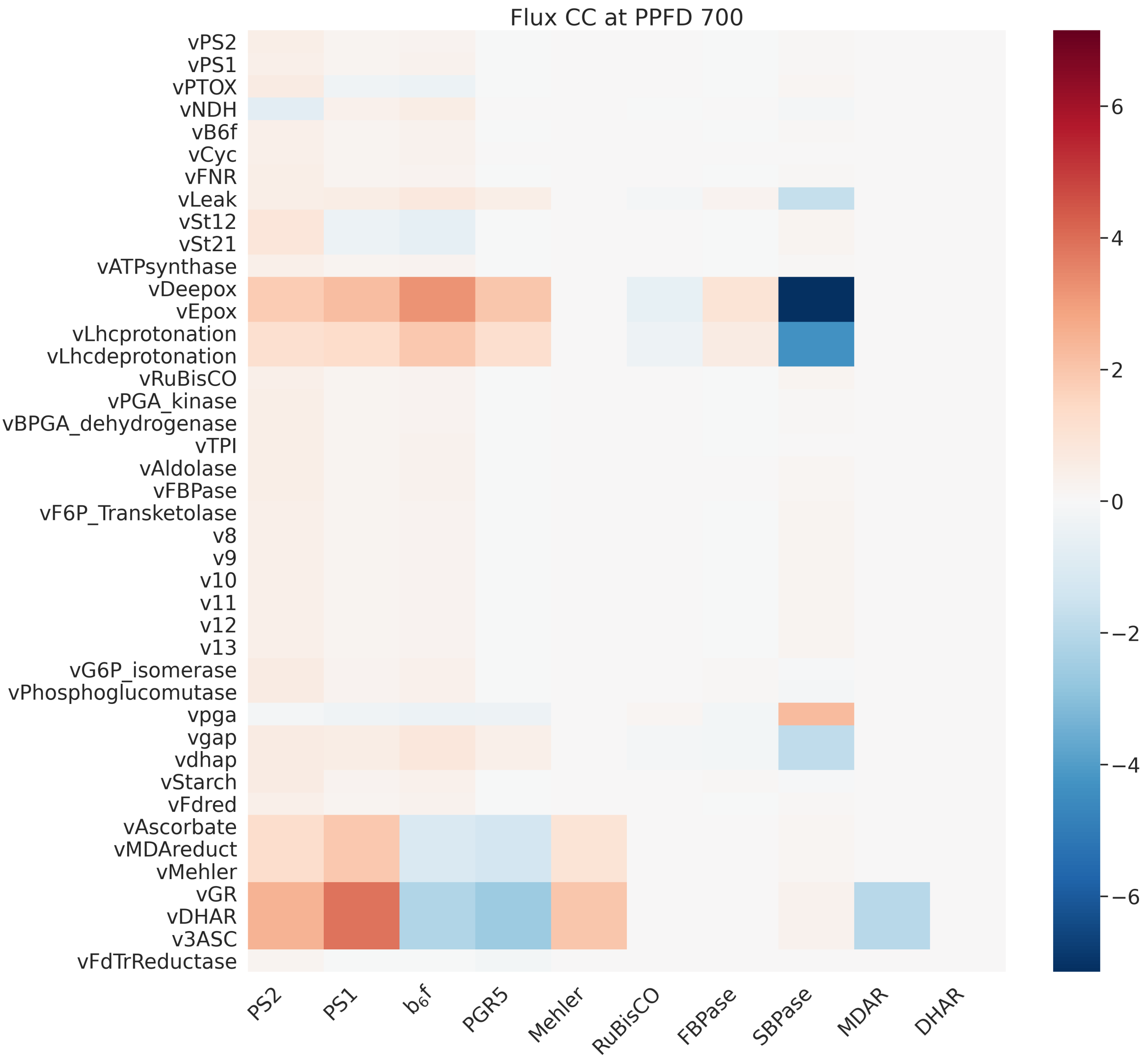


Figure S23: Flux control coefficients of representative reactions of photosynthesis at specified PPFD.

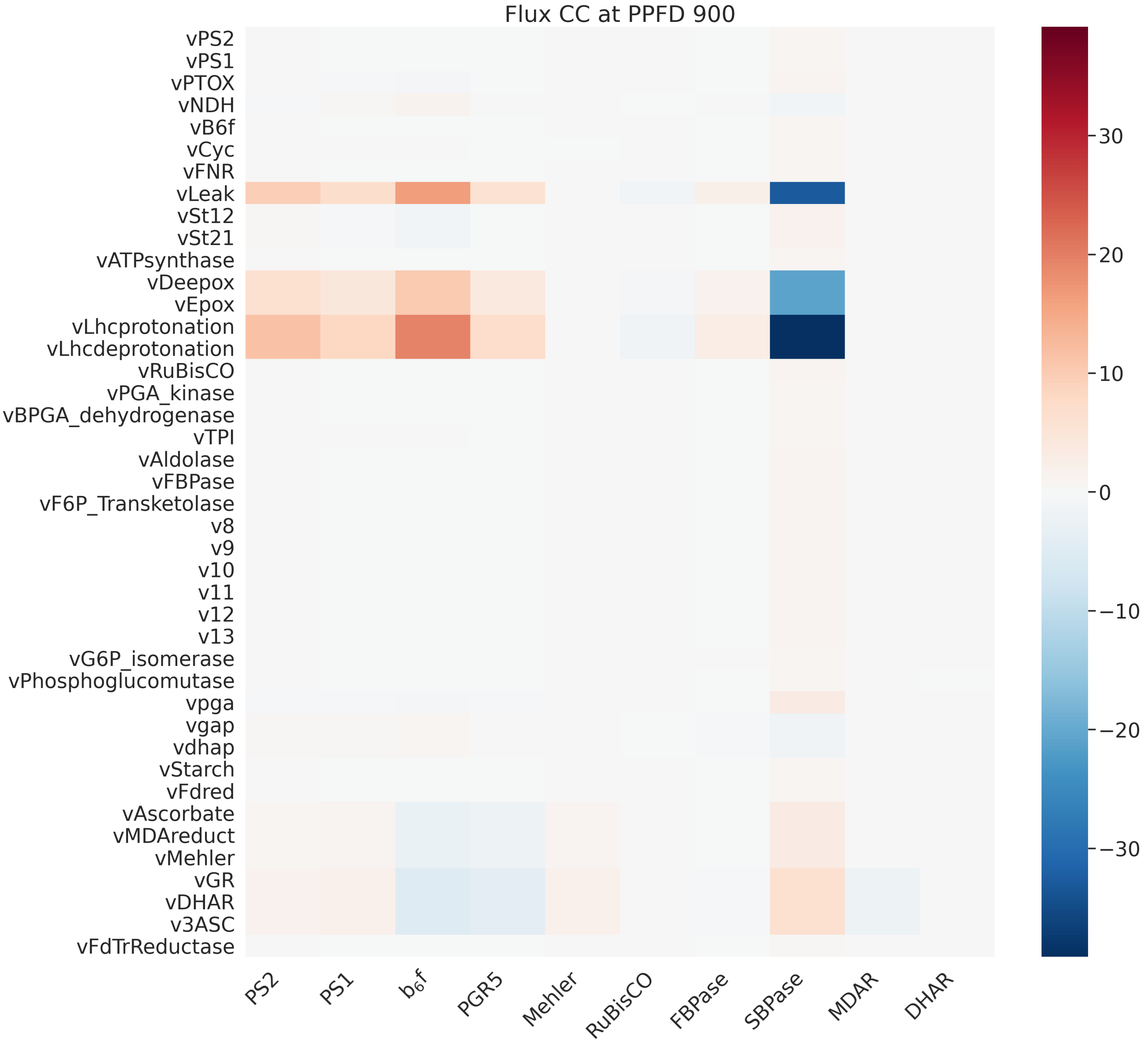


Figure S24: Flux control coefficients of representative reactions of photosynthesis at specified PPFD.

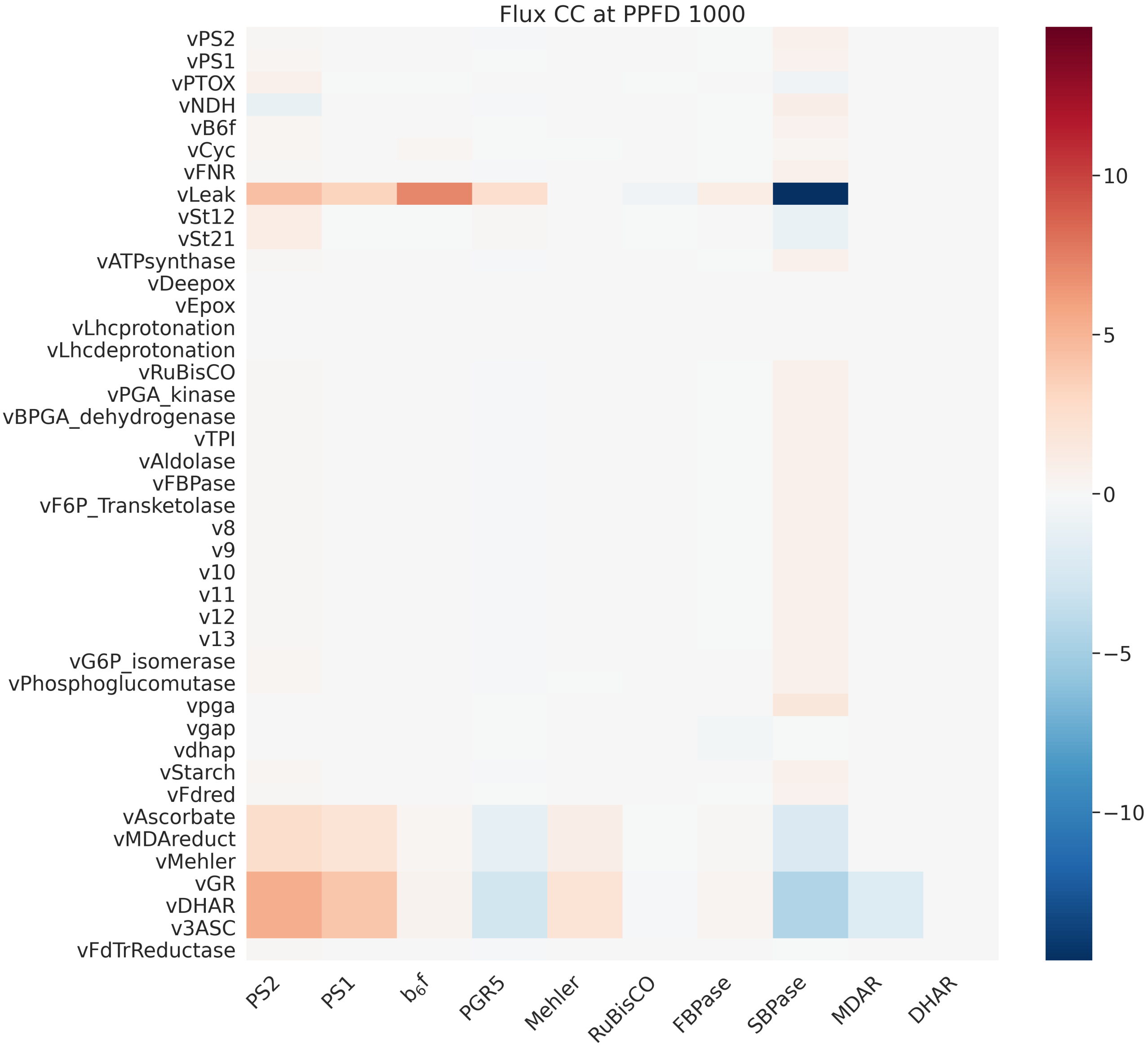


Figure S25: Flux control coefficients of representative reactions of photosynthesis at specified PPFD.

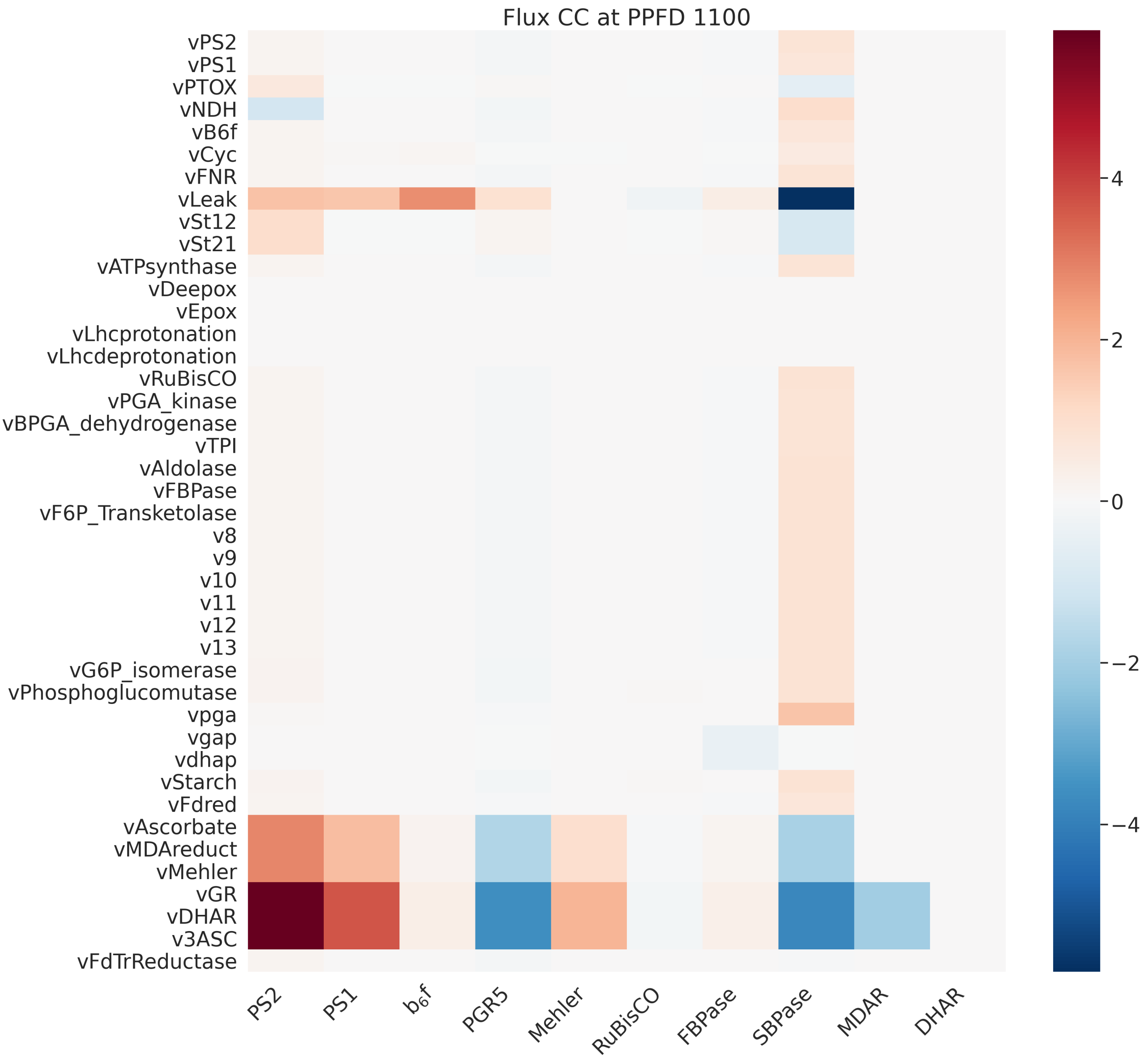


Figure S26: Flux control coefficients of representative reactions of photosynthesis at specified PPFD.

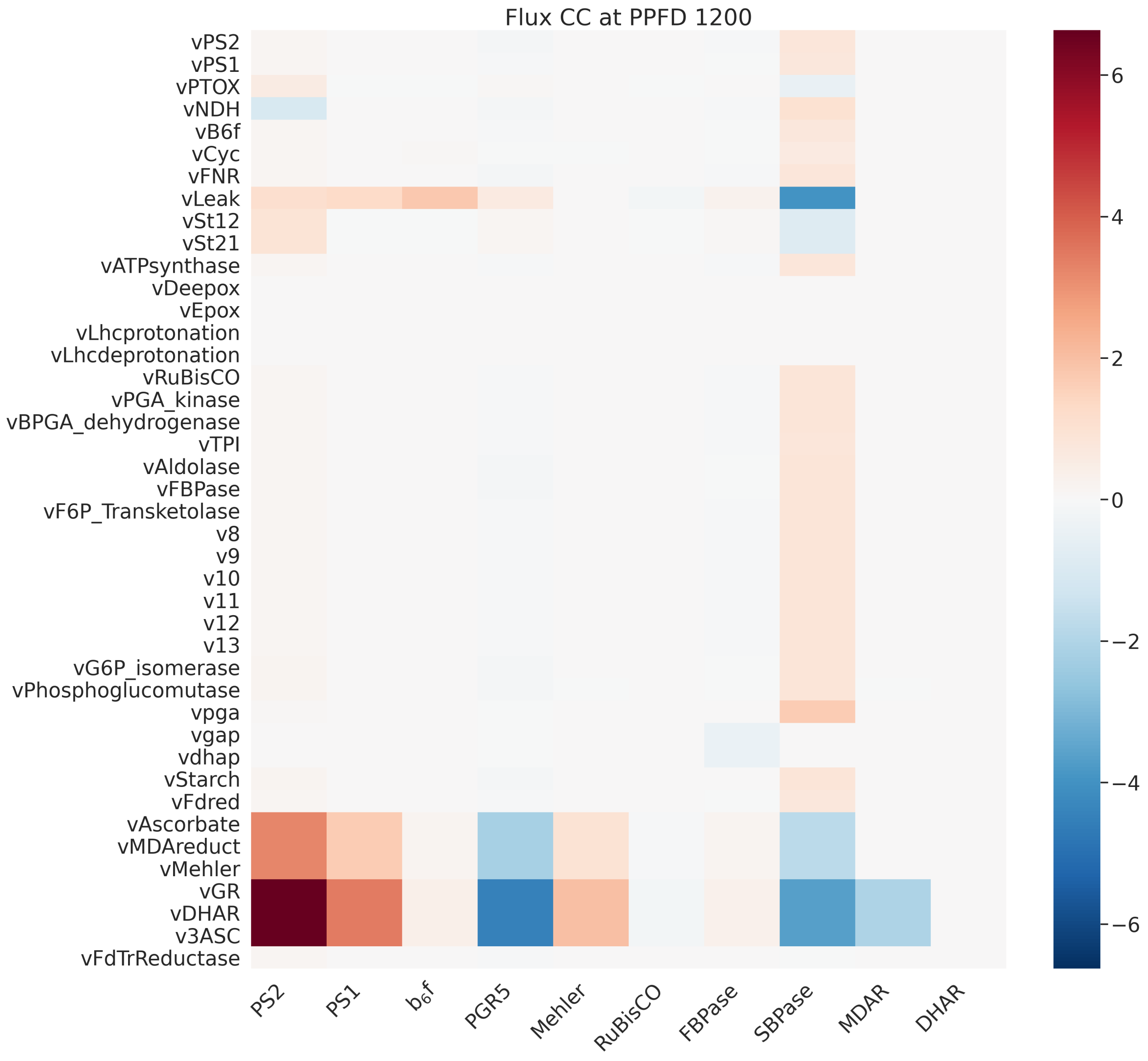


Figure S27: Flux control coefficients of representative reactions of photosynthesis at specified PPFD.

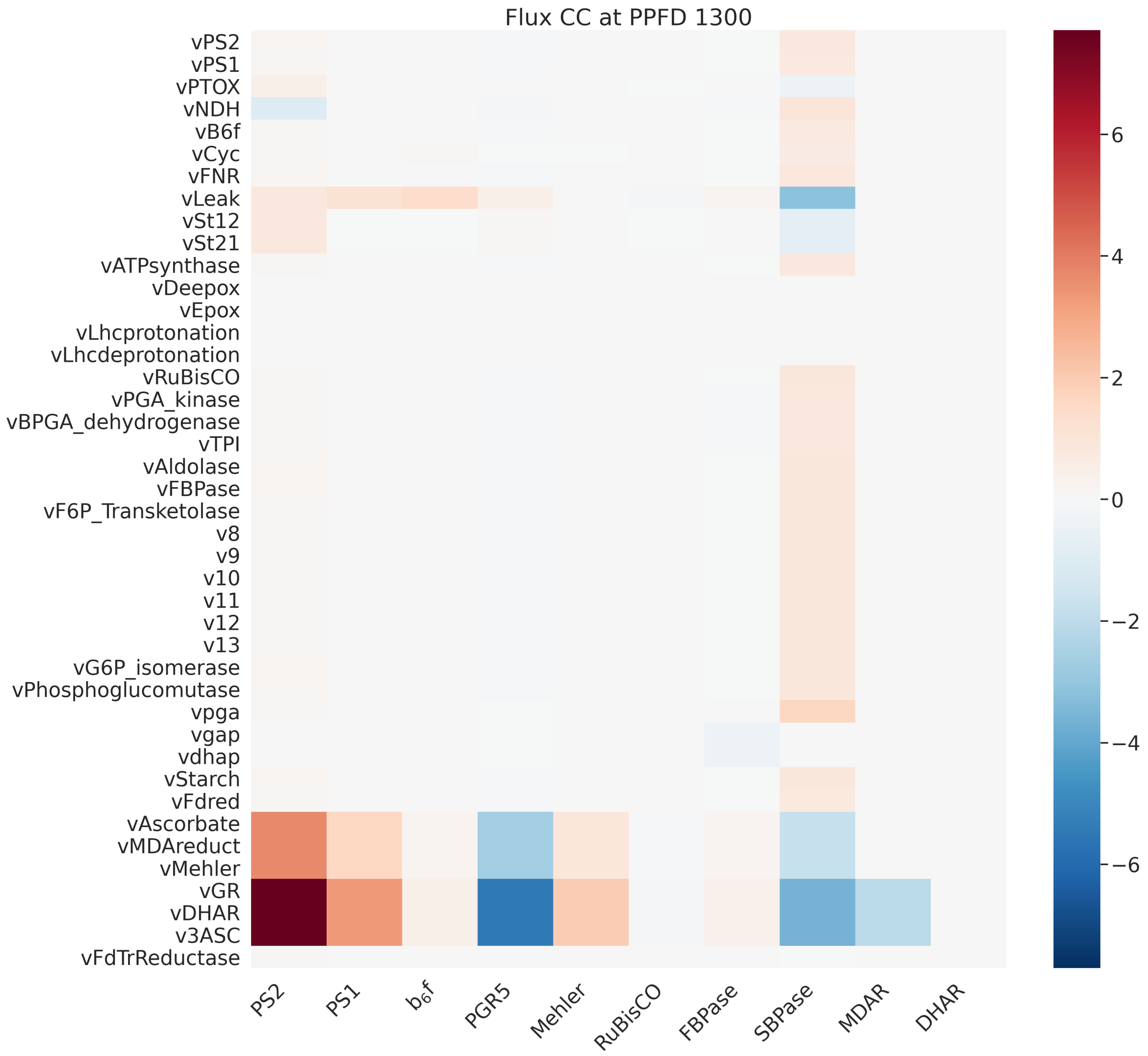


Figure S28: Flux control coefficients of representative reactions of photosynthesis at specified PPFD.

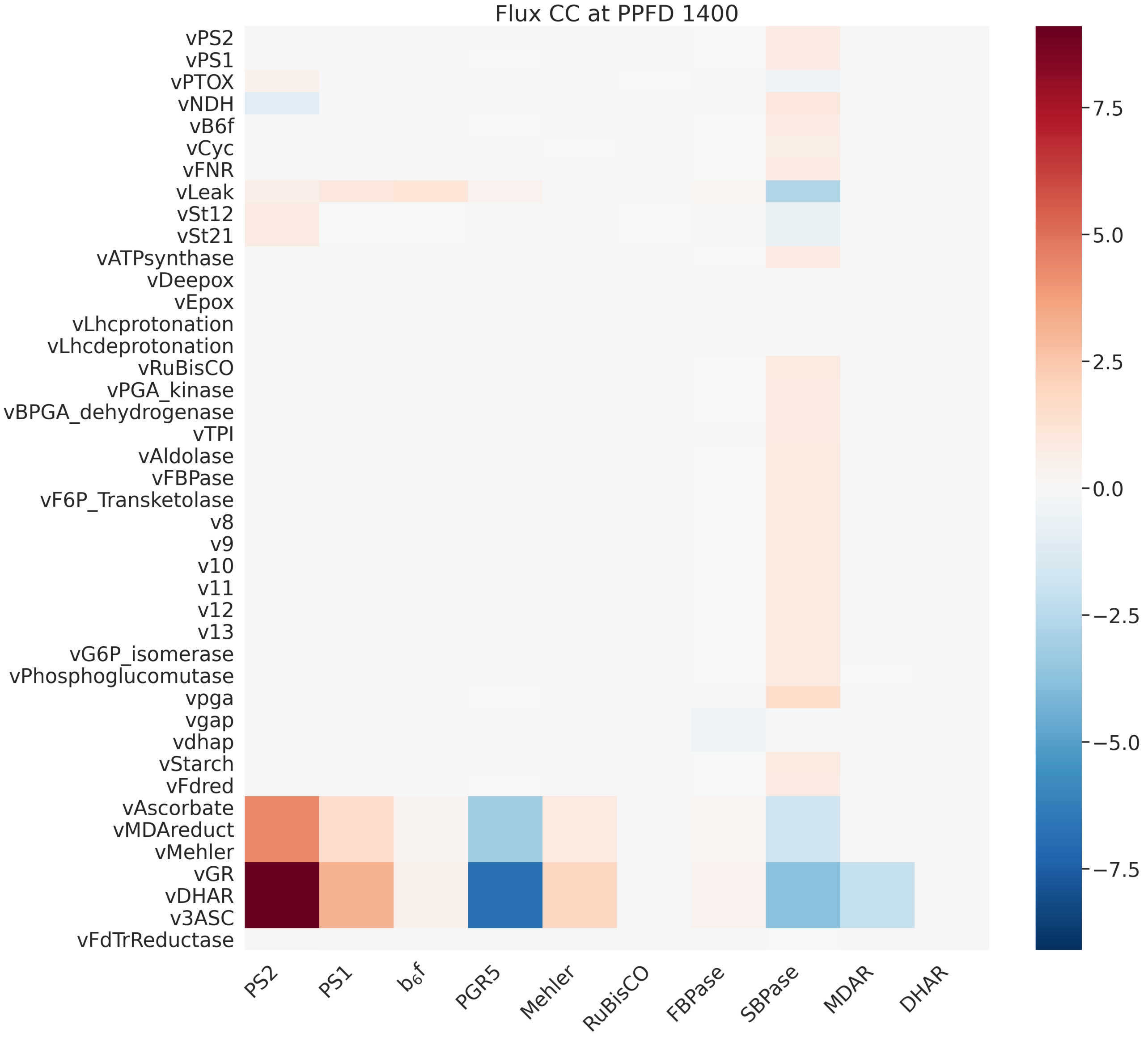


Figure S29: Flux control coefficients of representative reactions of photosynthesis at specified PPFD.

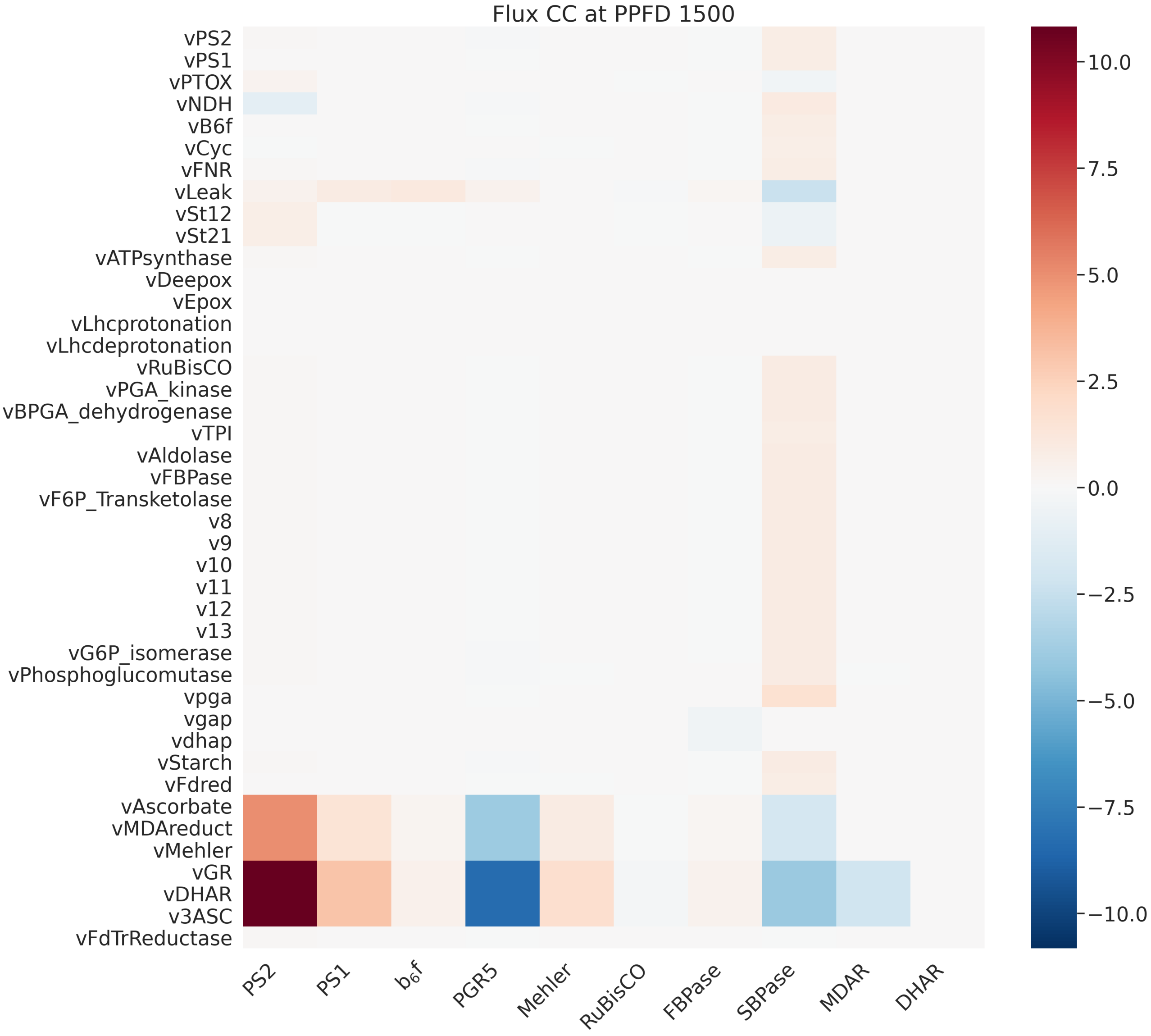


Figure S30: Flux control coefficients of representative reactions of photosynthesis at specified PPFD.

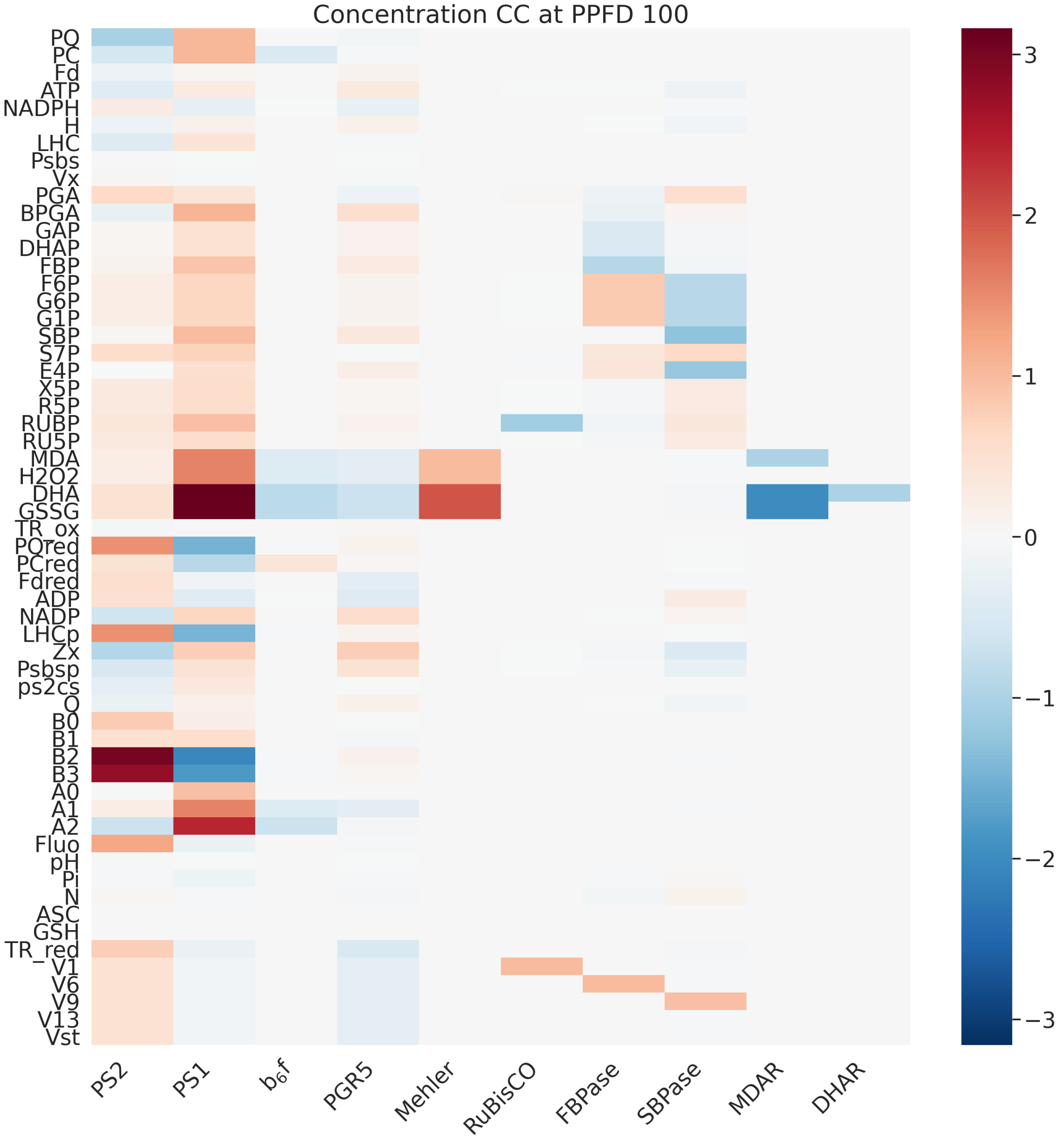


Figure S31: Concentration control coefficients of representative reactions of photosynthesis at specified PPFD.

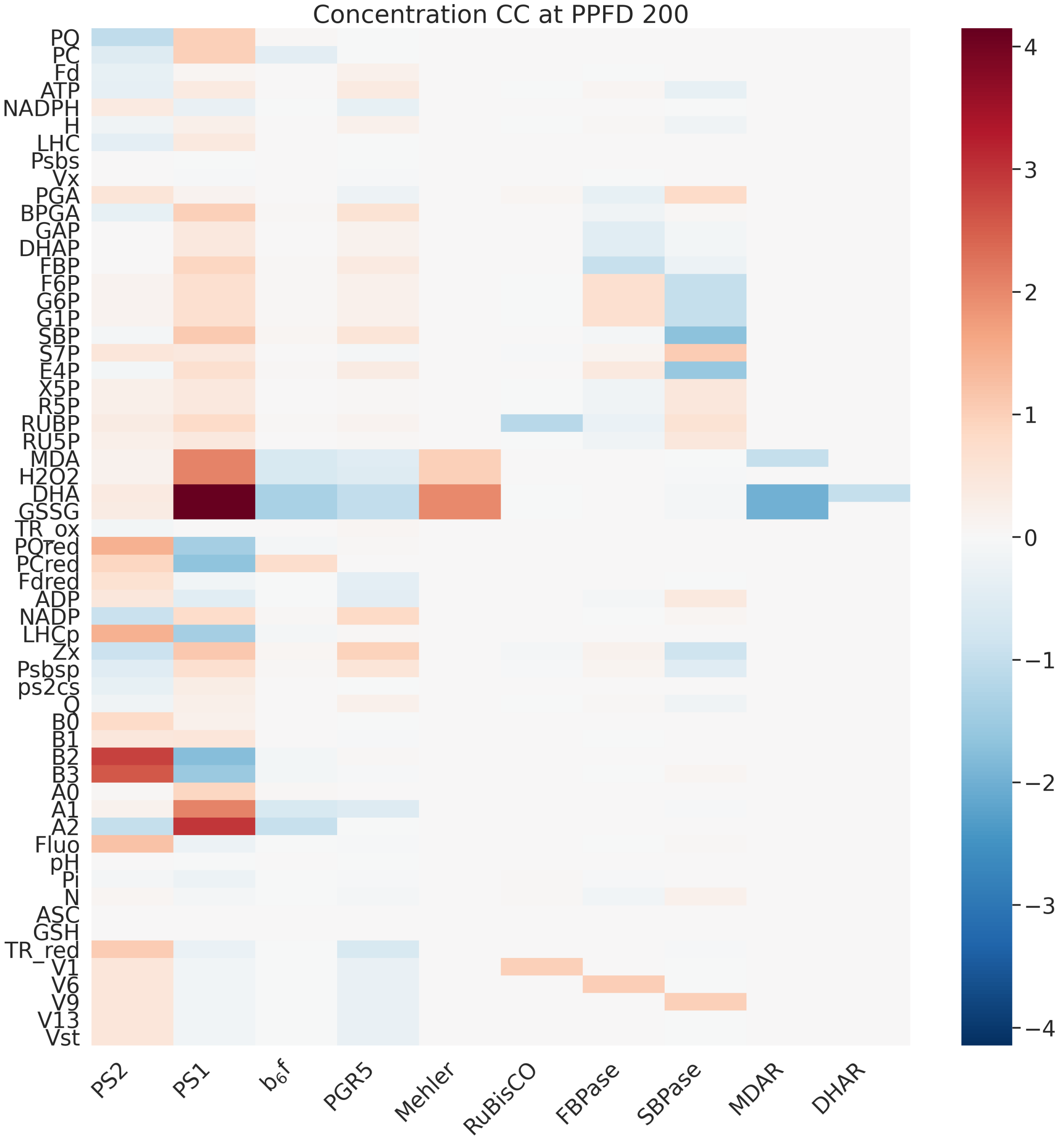


Figure S32: Concentration control coefficients of representative reactions of photosynthesis at specified PPFD.

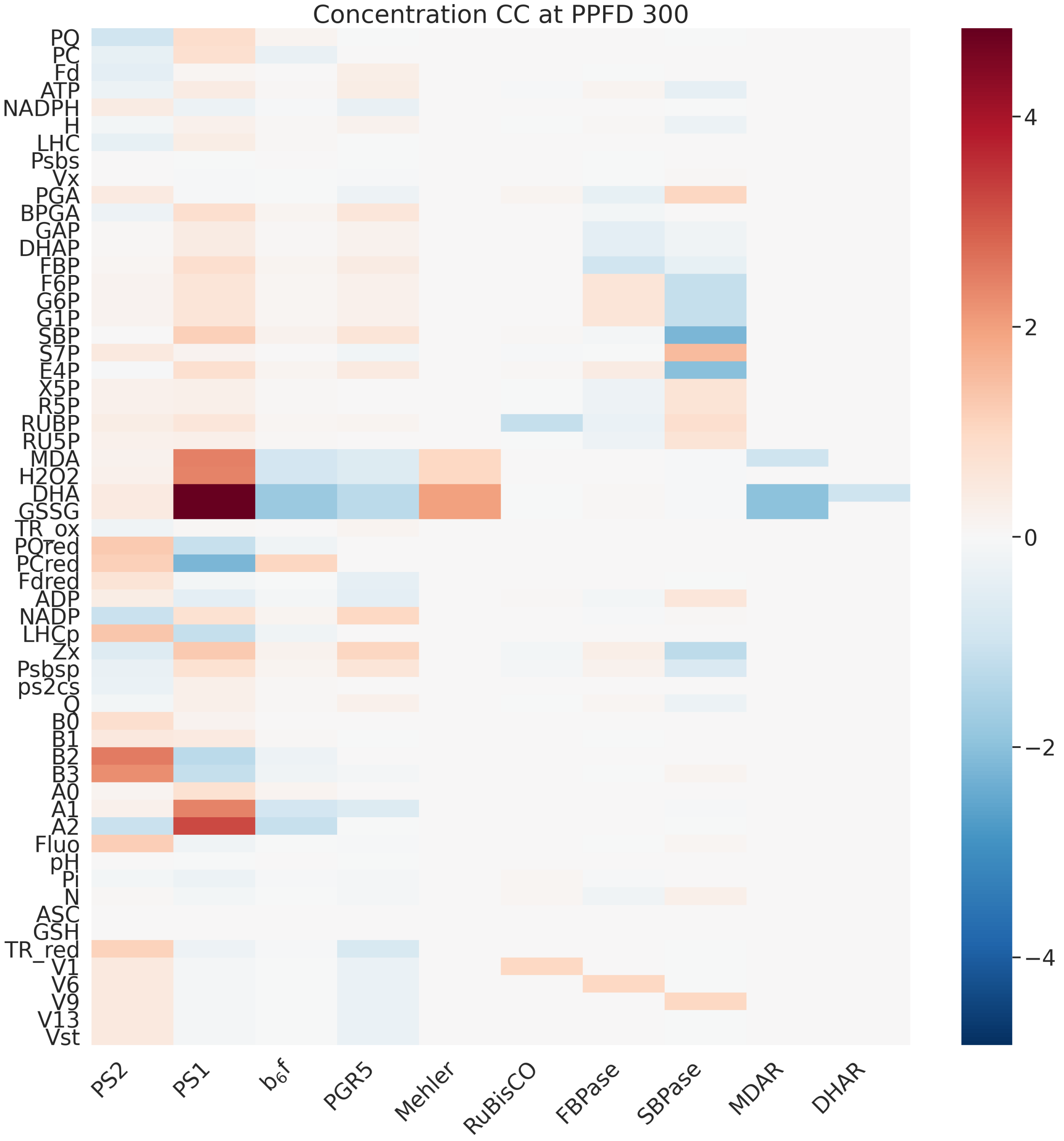


Figure S33: Concentration control coefficients of representative reactions of photosynthesis at specified PPFD.

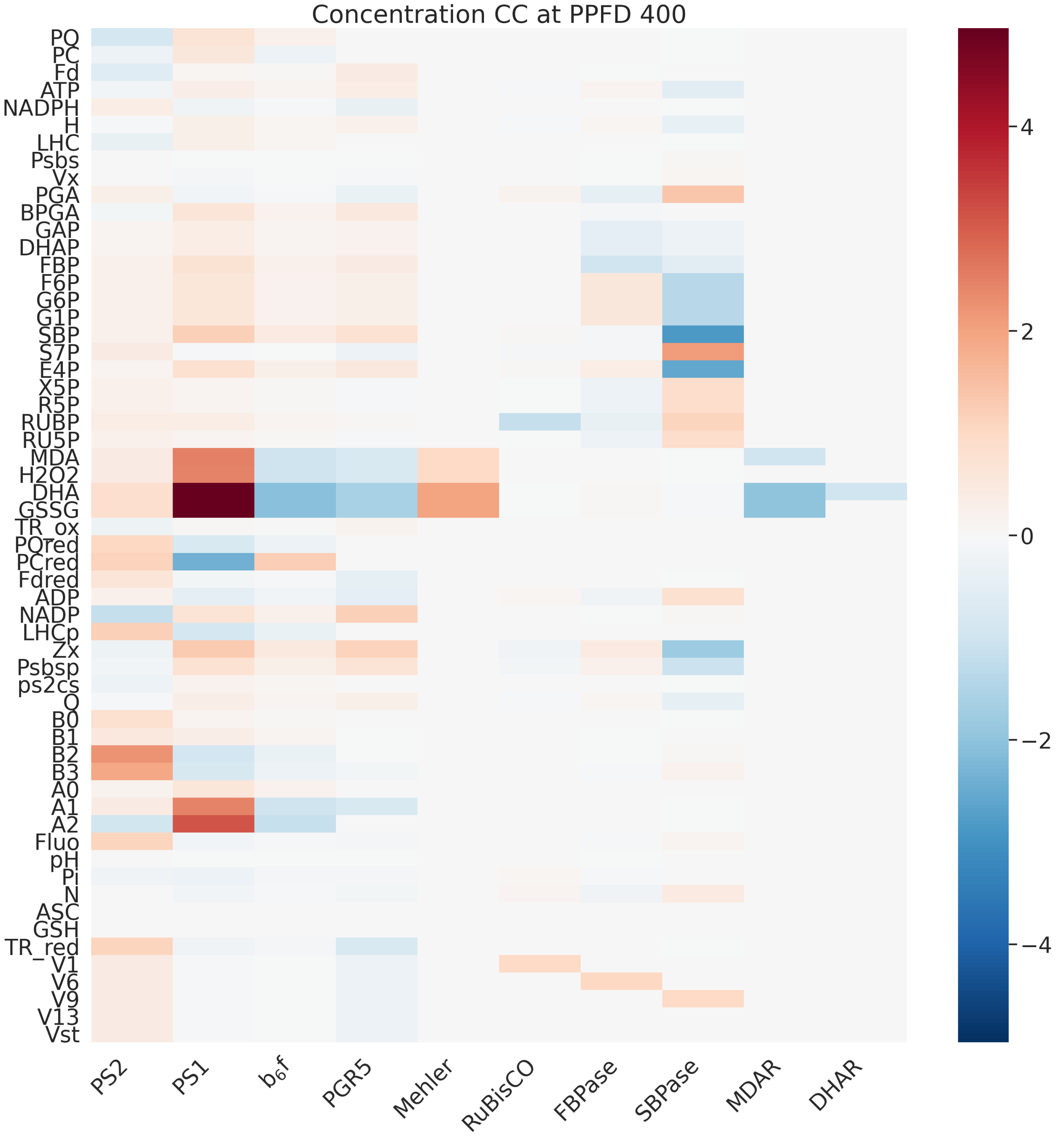


Figure S34: Concentration control coefficients of representative reactions of photosynthesis at specified PPFD.

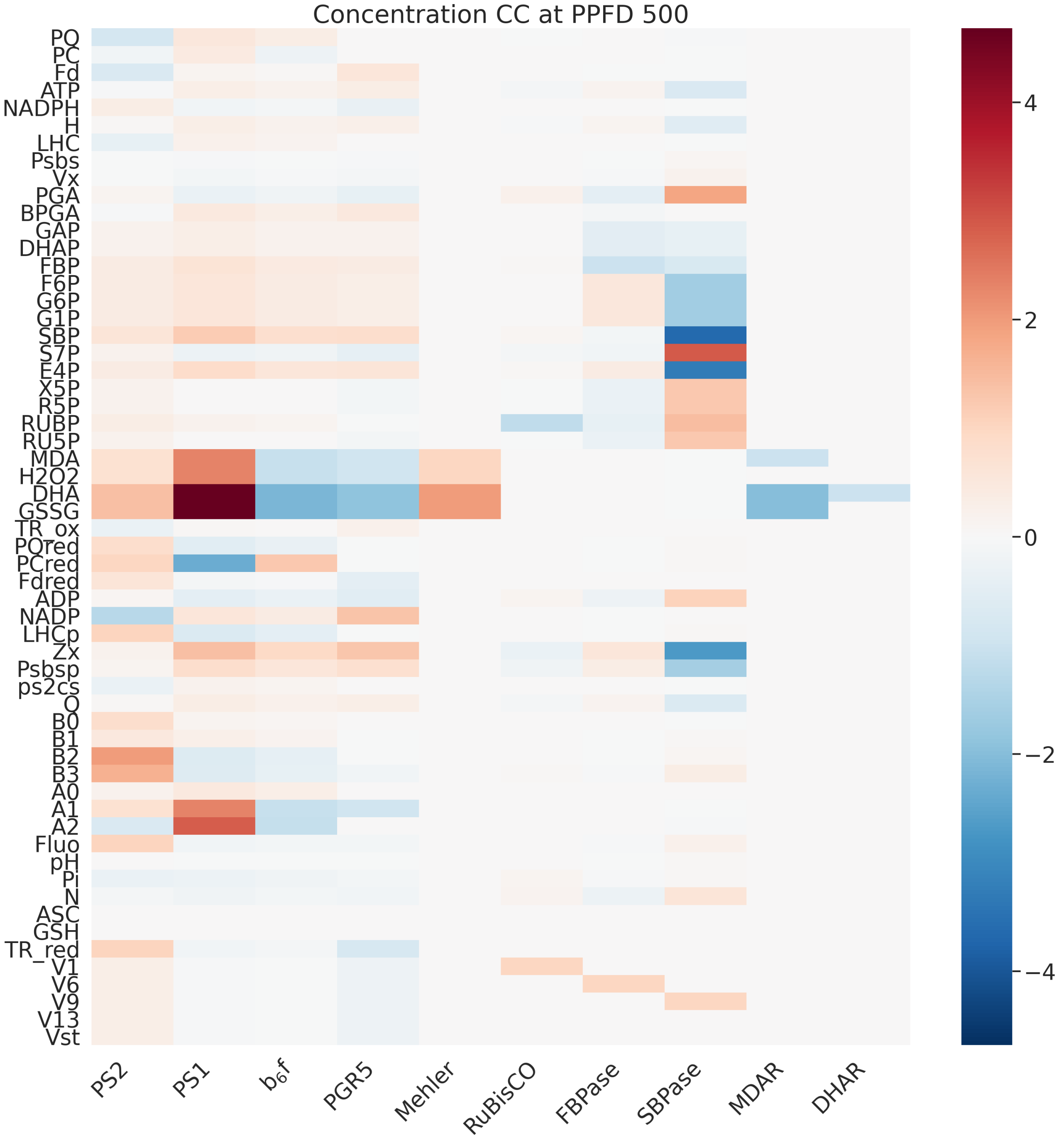


Figure S35: Concentration control coefficients of representative reactions of photosynthesis at specified PPFD.

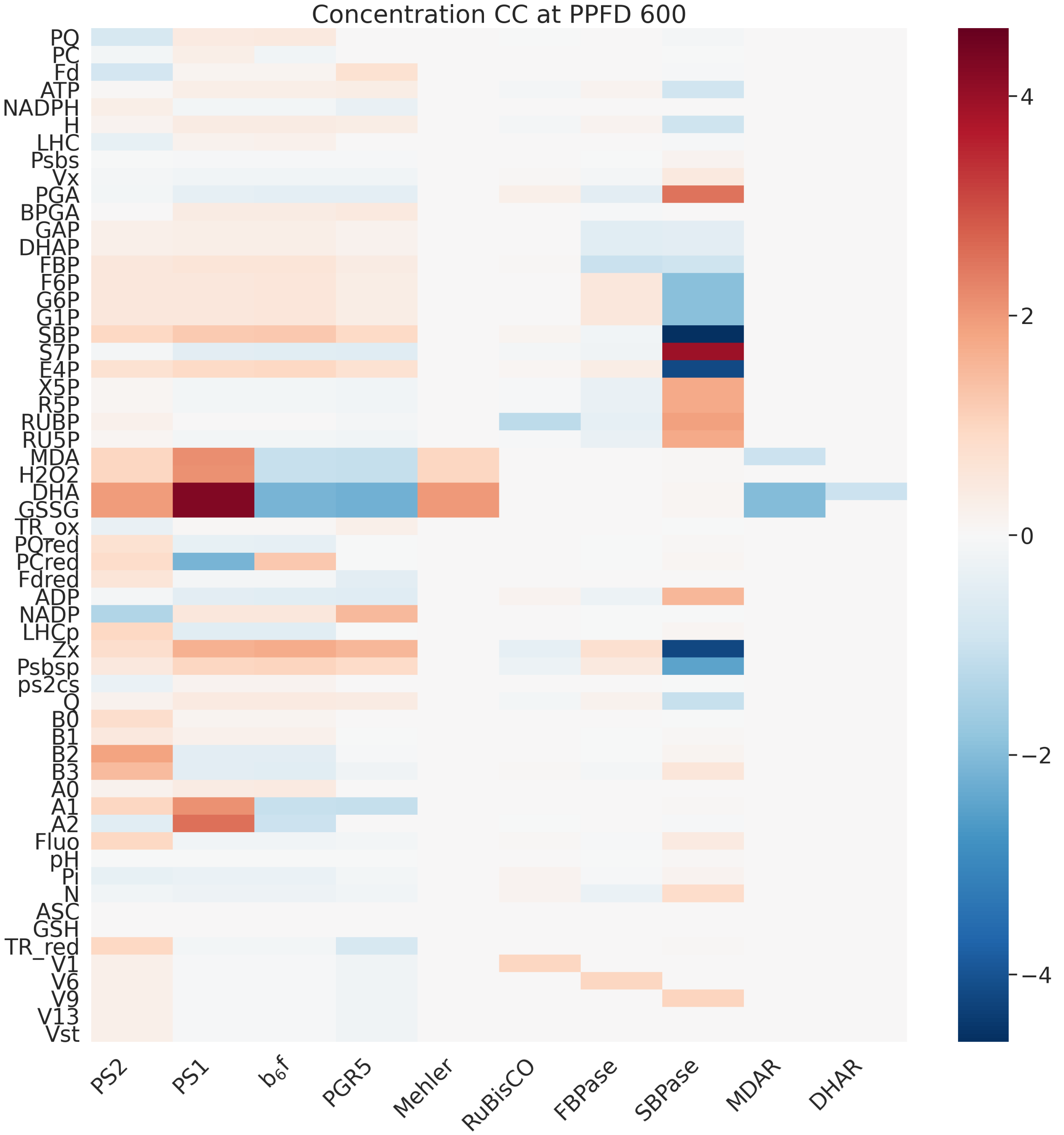


Figure S36: Concentration control coefficients of representative reactions of photosynthesis at specified PPFD.

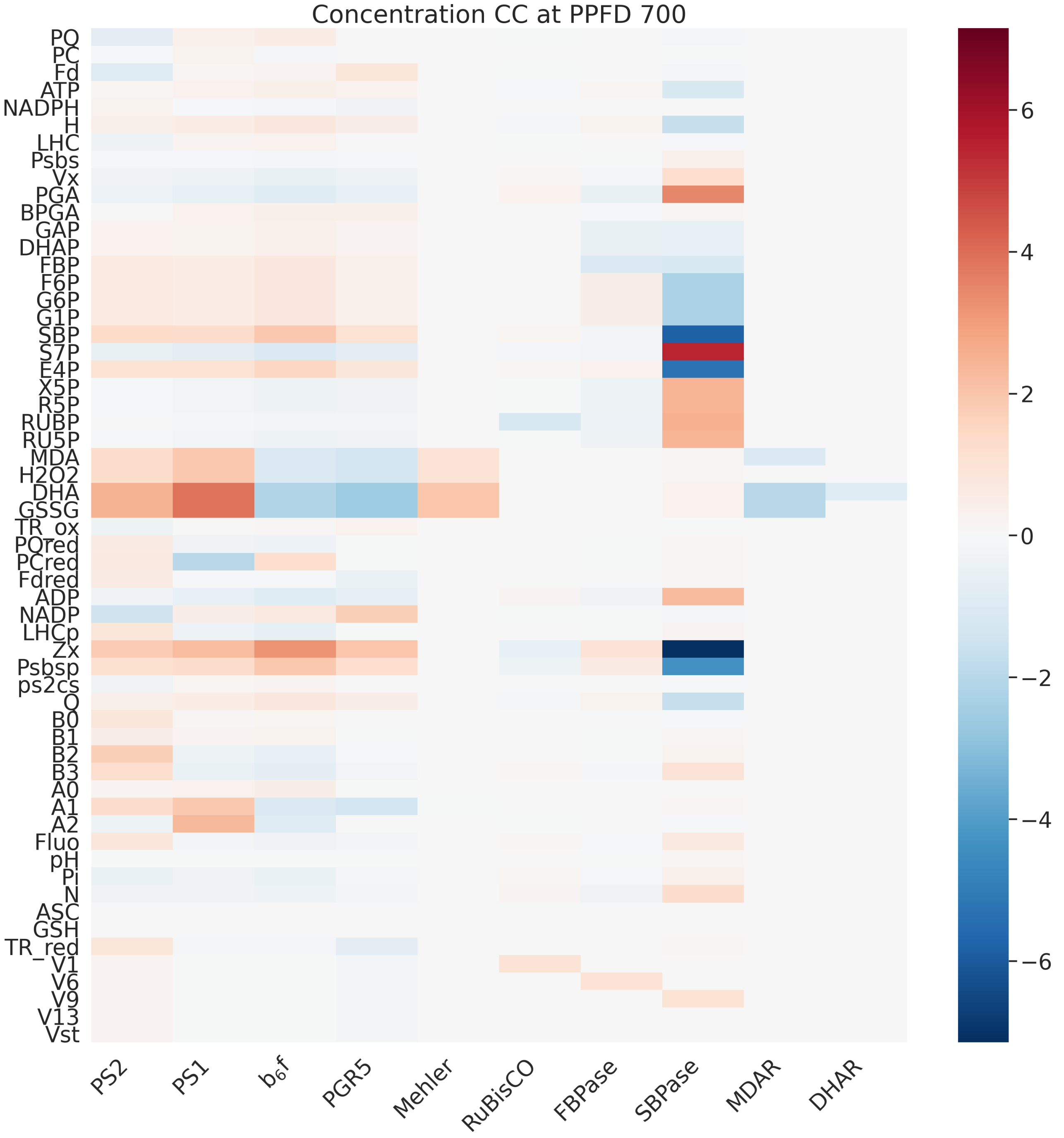


Figure S37: Concentration control coefficients of representative reactions of photosynthesis at specified PPFD.

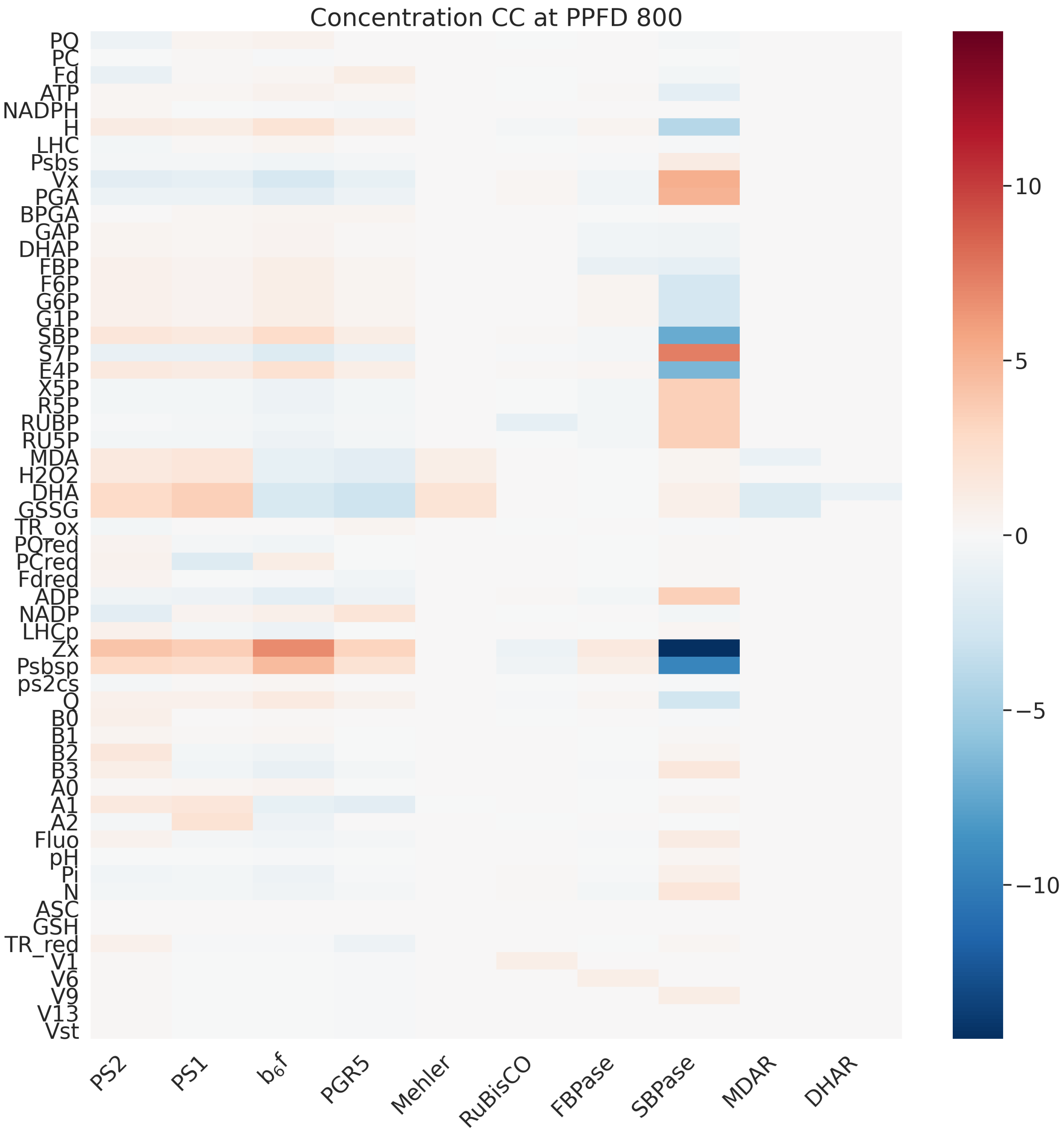


Figure S38: Concentration control coefficients of representative reactions of photosynthesis at specified PPFD.

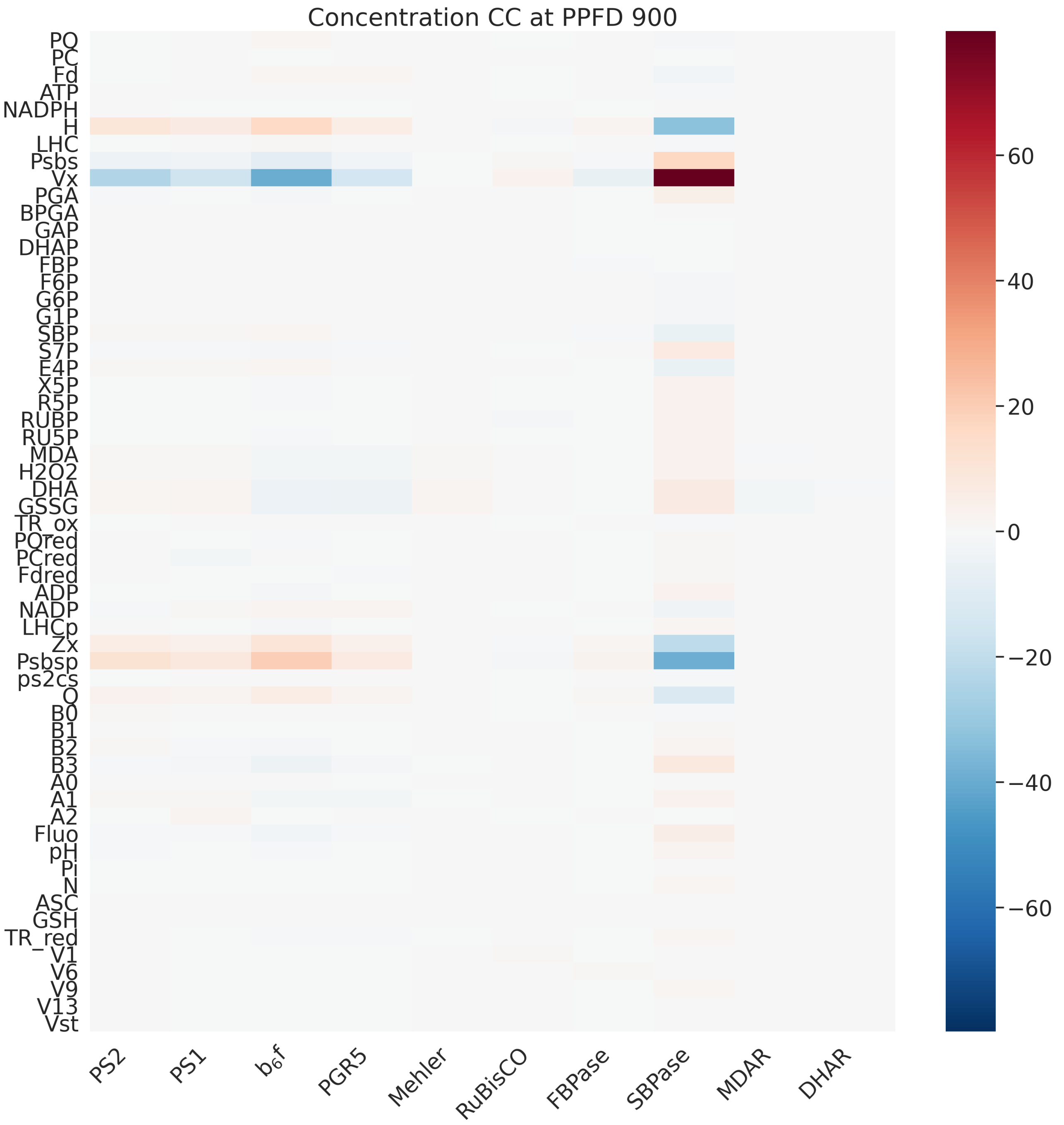


Figure S39: Concentration control coefficients of representative reactions of photosynthesis at specified PPFD.

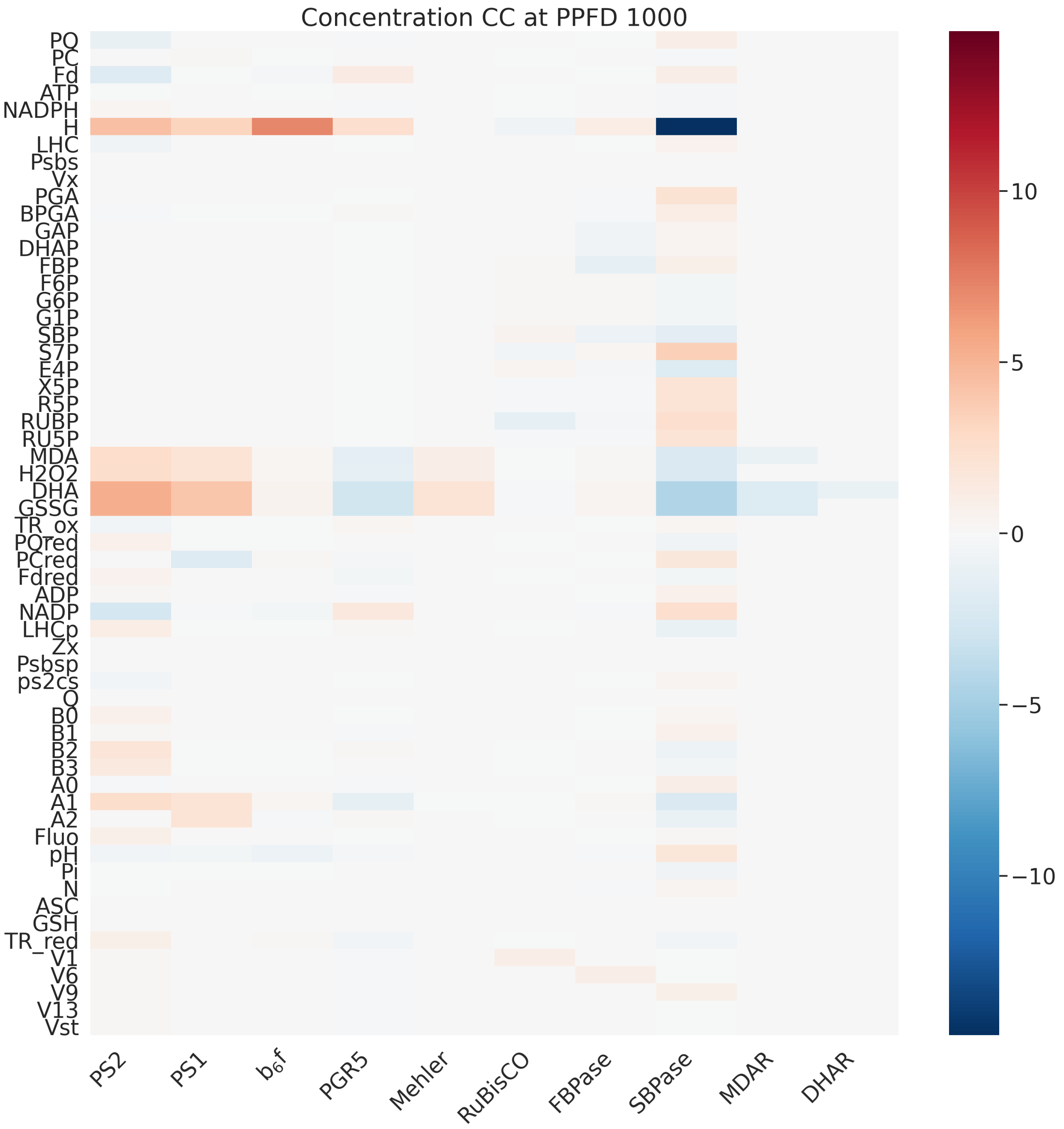


Figure S40: Concentration control coefficients of representative reactions of photosynthesis at specified PPFD.

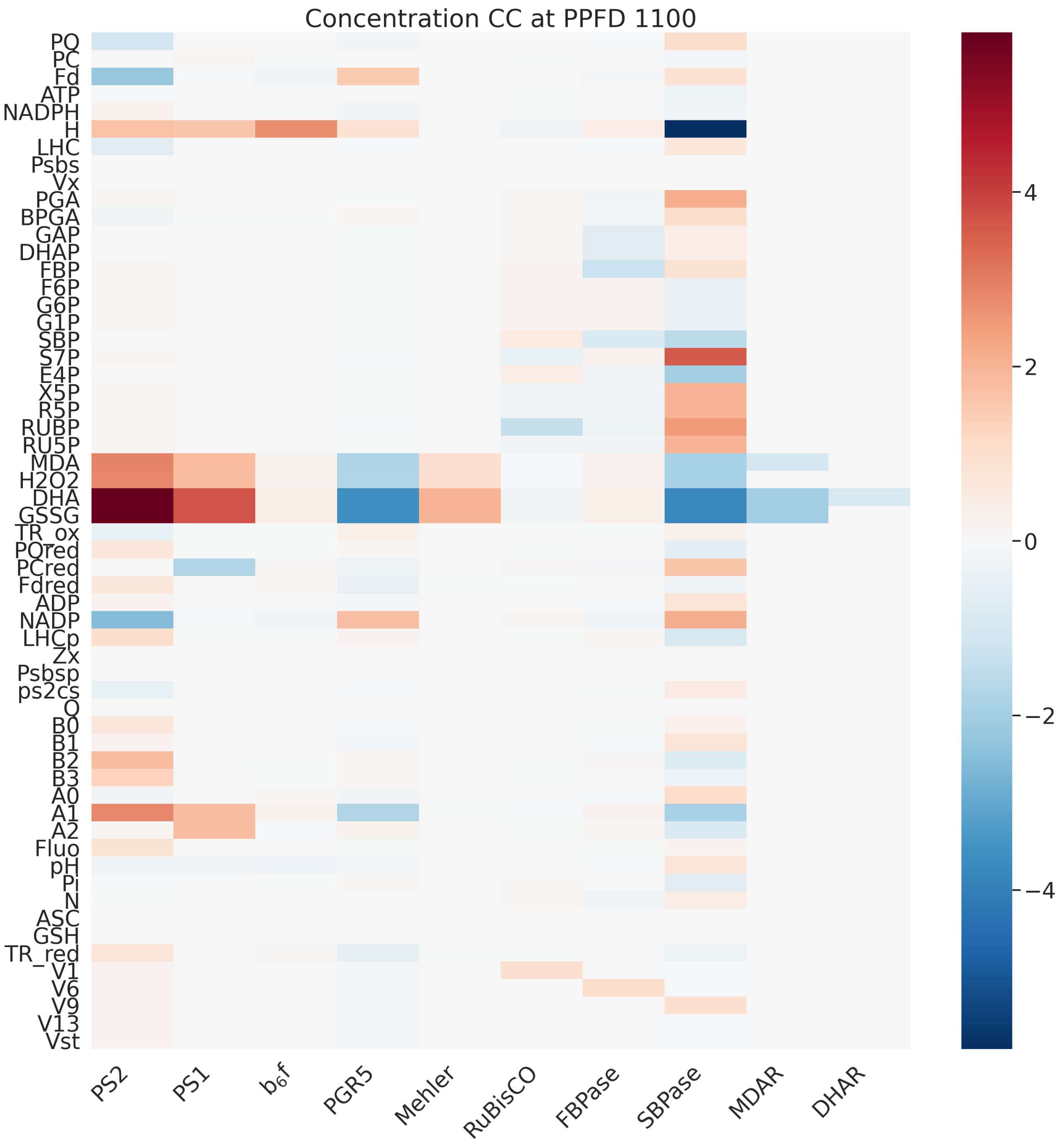


Figure S41: Concentration control coefficients of representative reactions of photosynthesis at specified PPFD.

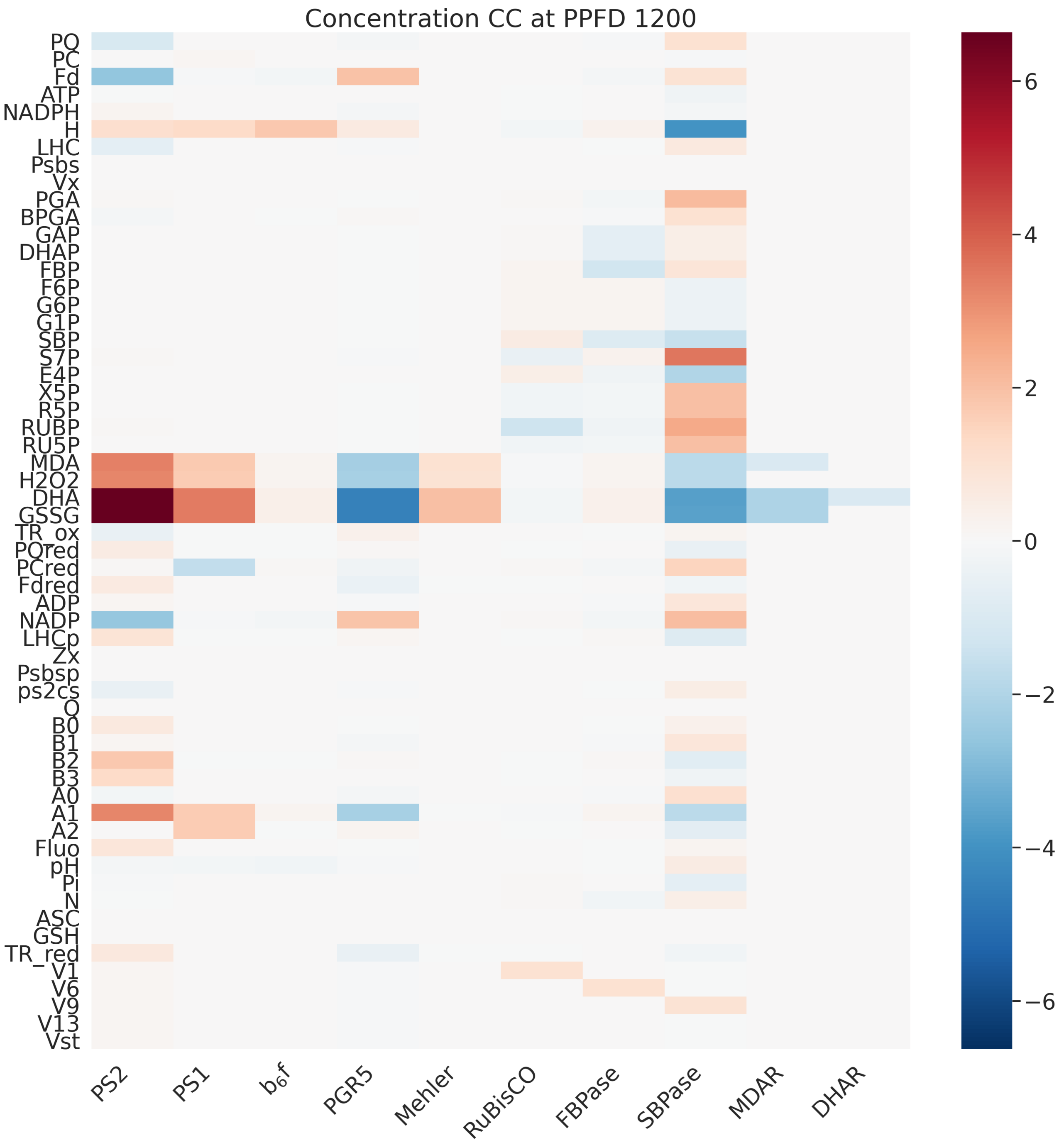


Figure S42: Concentration control coefficients of representative reactions of photosynthesis at specified PPFD.

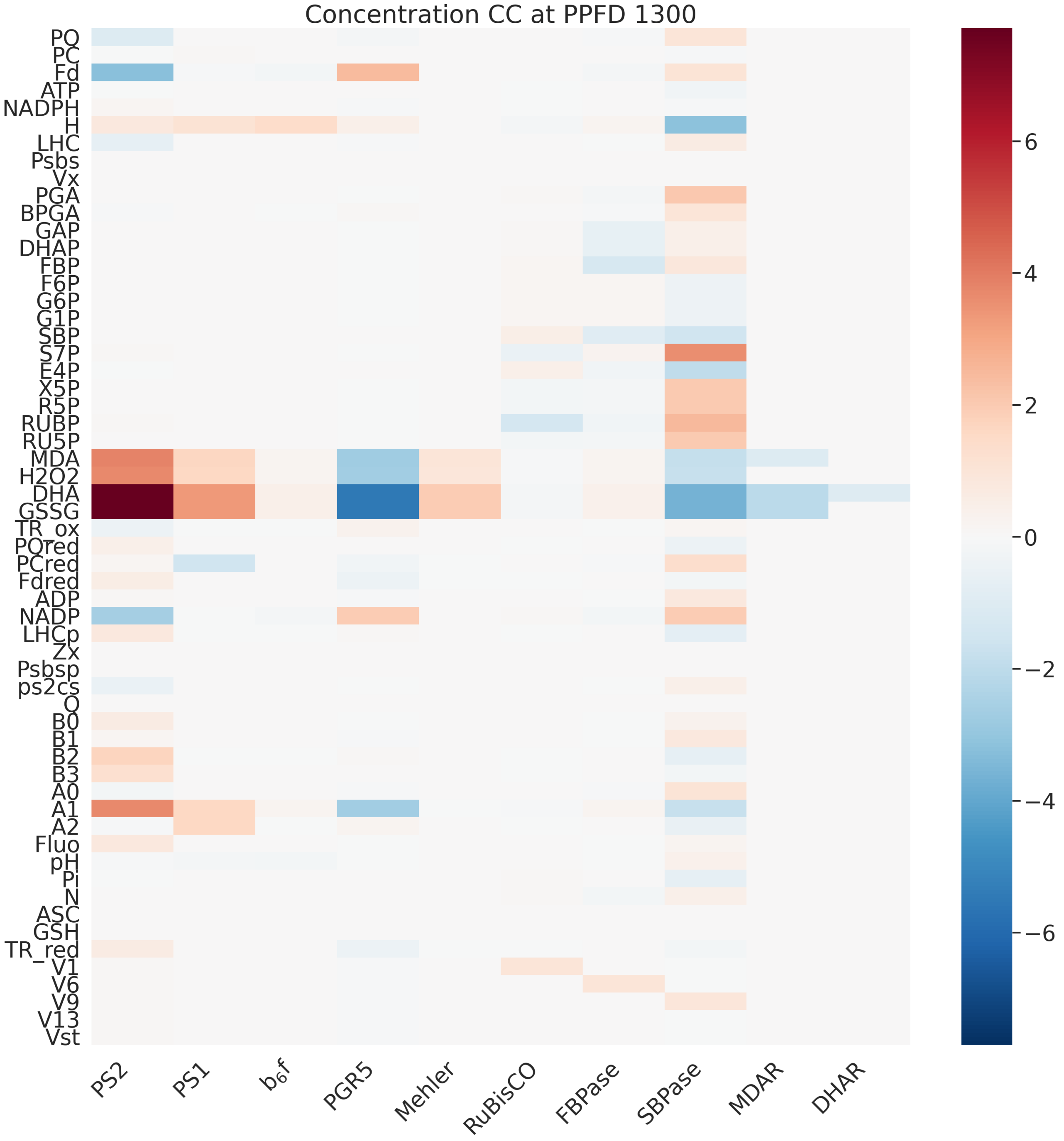


Figure S43: Concentration control coefficients of representative reactions of photosynthesis at specified PPFD.

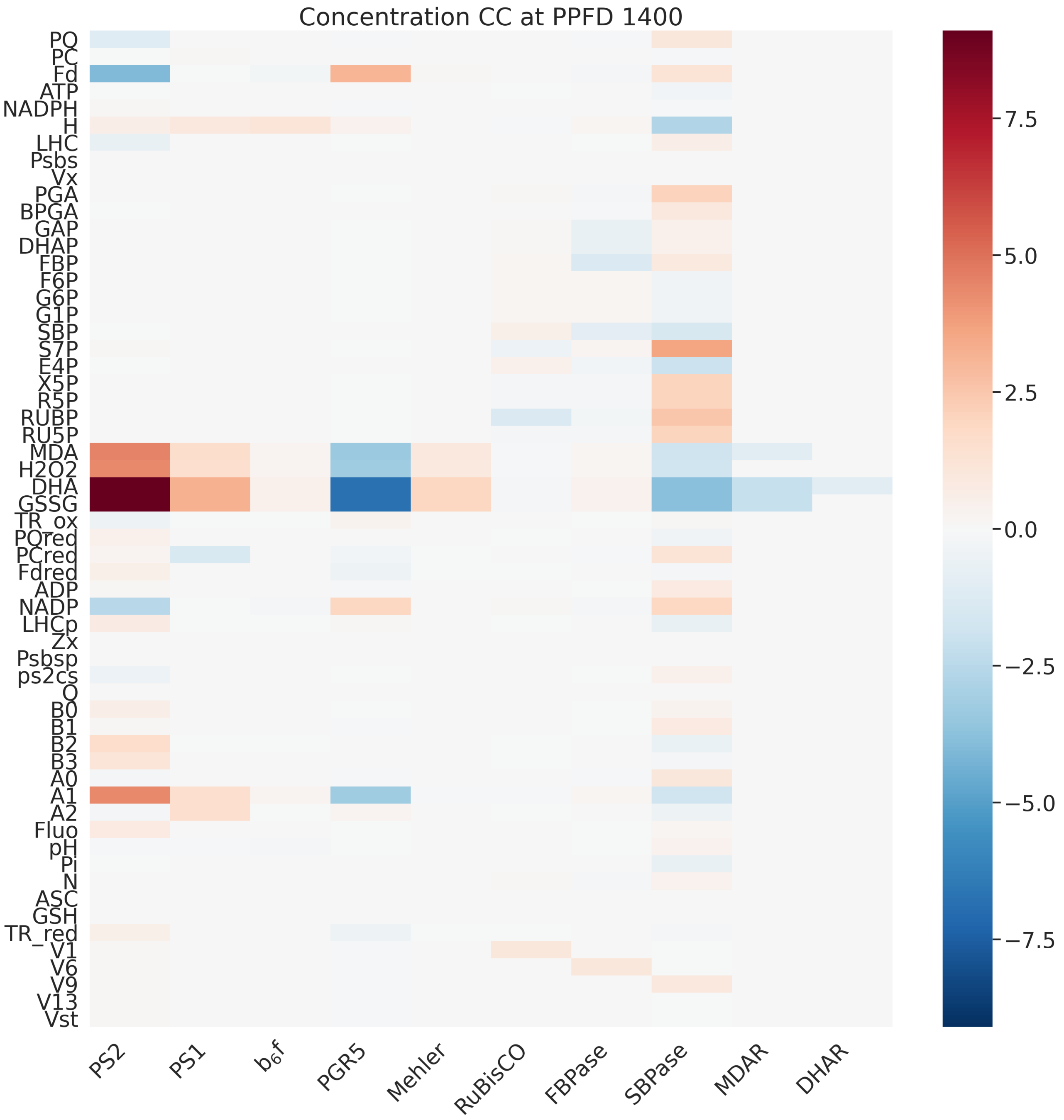


Figure S44: Concentration control coefficients of representative reactions of photosynthesis at specified PPFD.

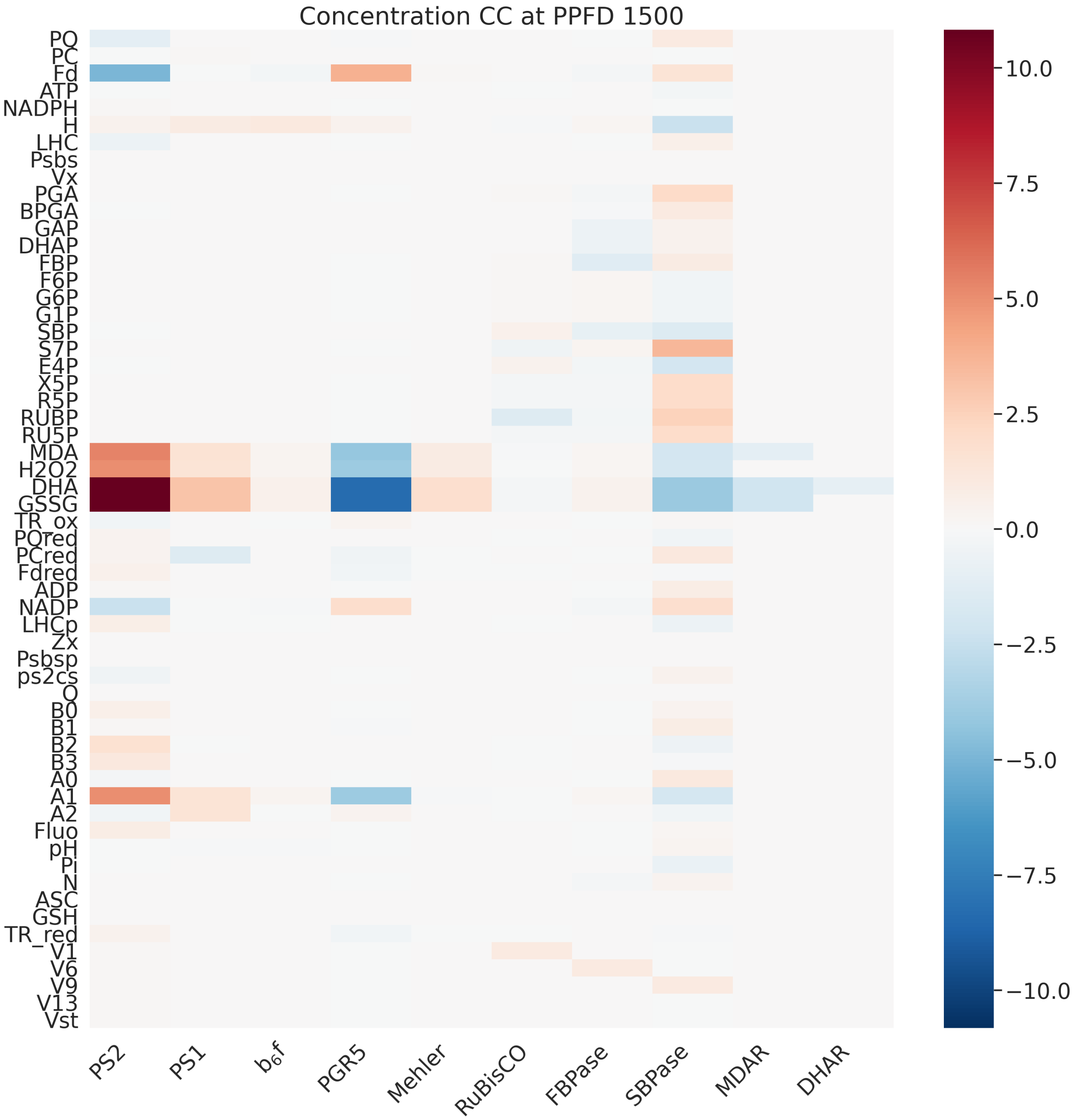


Figure S45: Concentration control coefficients of representative reactions of photosynthesis at specified PPFD.
